# Prime-seq, efficient and powerful bulk RNA-sequencing

**DOI:** 10.1101/2021.09.27.459575

**Authors:** Aleksandar Janjic, Lucas E. Wange, Johannes W. Bagnoli, Johanna Geuder, Phong Nguyen, Daniel Richter, Beate Vieth, Binje Vick, Irmela Jeremias, Christoph Ziegenhain, Ines Hellmann, Wolfgang Enard

## Abstract

With the advent of Next Generation Sequencing, RNA-sequencing (RNA-seq) has become the major method for quantitative gene expression analysis. Reducing library costs by early barcoding has propelled single-cell RNA-seq, but has not yet caught on for bulk RNA-seq. Here, we optimized and validated a bulk RNA-seq method we call prime-seq. We show that with respect to library complexity, measurement accuracy, and statistical power it performs equivalent to TruSeq, a standard bulk RNA-seq method, but is four-fold more cost-efficient due to almost 50-fold cheaper library costs. We also validate a direct RNA isolation step that further improves cost and time-efficiency, show that intronic reads are derived from RNA, validate that prime-seq performs optimal with only 1,000 cells as input, and calculate that prime-seq is the most cost-efficient bulk RNA-seq method currently available. We discuss why many labs would profit from a cost-efficient early barcoding RNA-seq protocol and argue that prime-seq is well suited for setting up such a protocol as it is well validated, well documented, and requires no specialized equipment.

## Background

RNA-sequencing (RNA-seq) has become a central method in biology and many technological variants exist that are adapted to different biological questions [1]. Its most frequent application is the quantification of gene expression levels to identify differentially expressed genes, infer regulatory networks, or identify cellular states. This is done on populations of cells (bulk RNA-seq) and increasingly with single-cell or single-nucleus resolution (scRNA-seq). Choosing a suitable RNA-seq method for a particular biological question depends on many aspects, but the number of samples that can be analyzed is almost always a crucial factor. Including more biological replicates increases the power to detect differences and including more sample conditions increases the generalizability of the study. As the limiting factor for the number of samples is often the budget, the costs of an RNA-seq method are an essential parameter for the biological insights that can be gained from a study. Of note, costs need to be viewed in the context of statistical power, i.e. in light of the true and false positive rate of a method [2,3] and these “normalized” costs can be seen as cost-efficiency. On top of reagent costs per sample, aspects like robustness, hands-on time, and setup investments of a method can also be seen as cost factors. Other important factors less directly related to cost efficiency are the number and types of genes that can be detected (complexity), the amount of input material that is needed to detect them (sensitivity), and how well the measured signal reflects the actual transcript concentration (accuracy).

In recent years, technological developments have focused on scRNA-seq due to its exciting possibilities and due to the urgent need to improve its cost efficiency and sensitivity [4–6]. A decisive development for cost efficiency was “early-barcoding”, i.e. the integration of sample-specific DNA tags in the primers used during complementary DNA (cDNA) generation [7,8]. This allows one to pool cDNA for all further library preparation steps, saving time and reagents. However, the cDNA and the barcode need to be sequenced from the same molecule and hence cDNA-tags and not full-length cDNA sequences are generated. An improvement in measurement noise is achieved by integrating a random DNA tag along with the sample barcode, a Unique Molecular Identifier (UMI), that allows identifying PCR duplicates and is especially relevant for the small starting amounts in scRNA-seq [2,7,9]. Optimizing reagents and reaction conditions (e.g. [10,11] and the efficient generation of small reaction chambers such as microdroplets [12–14], further improved cost efficiency and sensitivity and resulted in the current standard of scRNA-seq, commercialized by 10X Genomics [5].

Despite these exciting developments, bulk RNA-seq is still widely used and – more importantly – still widely useful as it allows for more flexibility in the experimental design that can be advantageous and complementary to scRNA-seq approaches. For example, investigated cell populations might be homogenous enough to justify averaging, single-cell or single-nuclei suspensions might be difficult or impossible to generate, or single-cell or single-nucleus suspension might be biased towards certain cell types. Most trivial, but maybe most crucial, the number of replicates and conditions is limited due to the high costs of scRNA-seq per sample. Furthemore, as more knowledge on cellular and spatial heterogeneity is acquired by scRNA-seq and spatial approaches, bulk RNA-seq profiles can be better interpreted, e.g. by computational deconvolution of the bulk profile [15]. Hence, bulk RNA-seq will remain a central method in biology, despite or even because of the impressive developments from scRNA-seq and spatial transcriptomics. However, bulk RNA-seq libraries are still largely made by isolating and fragmenting mRNA to generate random primed cDNA sequencing libraries. Commercial variants of such protocols, such as TruSeq and NEBNext, can be considered the current standard for bulk RNA-seq methods. This is partly because improvements of sensitivity and cost efficiency were less urgent for bulk RNA-seq as input amounts were often high, overall expenses were dominated by sequencing costs, and n=3 experimental designs have a long tradition in experimental biology [16]. However, input amounts can be a limiting factor, sequencing costs have decreased and will further decrease, and low sample size is a central problem of reproducibility [17,18]. To address these needs, several protocols have been developed, including targeted approaches [19–21] and genome-wide approaches that leverage the scRNA-seq developments described above [16,22]. However, given the importance and costs of bulk RNA-seq, even seemingly small changes, e.g. in the sequencing design of libraries [16], the number of PCR cycles [9], or enzymatic reactions [22], can have relevant impacts on cost efficiency, complexity, accuracy, and sensitivity. Furthermore, protocols need to be available to many labs to be useful and insufficient documentation, limited validation, and/or setup costs can prevent their implementation. Accordingly, further developments of bulk RNA-seq protocols are still useful.

Here, we have optimized and validated a bulk RNA-seq method that combines several methodological developments from scRNA-seq to generate a very sensitive and cost-efficient bulk RNA-seq method we call prime-seq (Figure 1, Figure S1). In particular, we have integrated and benchmarked a direct lysis and RNA purification step, validated that intronic reads are informative as they are not derived from genomic DNA, and show that prime-seq libraries are similar in complexity and statistical power to TruSeq libraries, but at least four-fold more cost-efficient due to almost 50-fold cheaper library costs. Prime-seq is also robust, as we have used variants of it in 22 publications [9,23–43], 132 experiments, and in 17 different organisms (Table S1, Figure S2). Additionally it has low setup costs as it does not require specialized equipment and is well validated and documented. Hence, it will be a very useful protocol for many labs or core facilities that quantify gene expression levels on a regular basis and have no cost-efficient protocol available yet.

**Figure 1.**
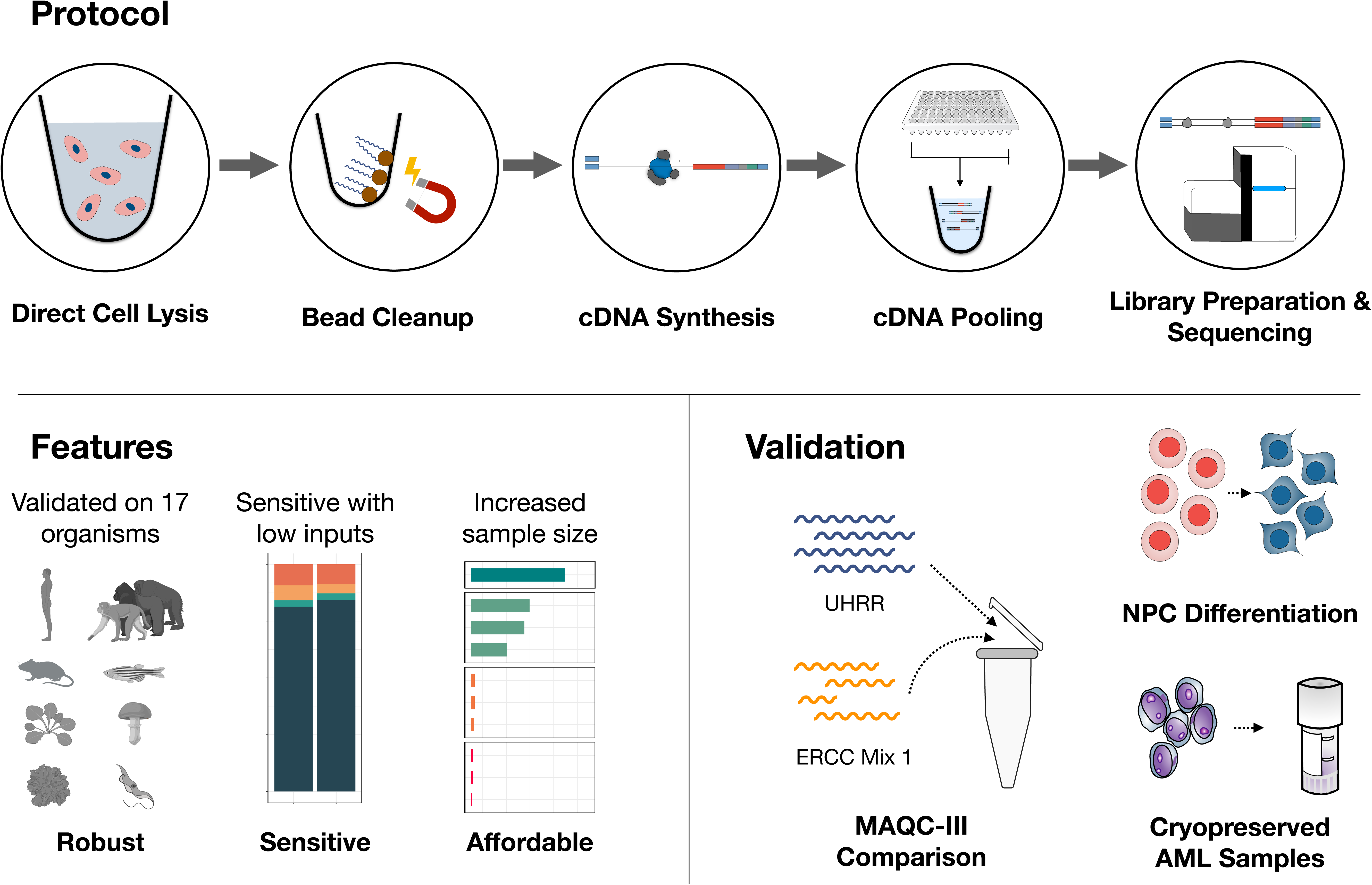
Graphical overview of prime-seq, highlighting its robustness, sensitivity, affordability, and the validation experiments performed. Cells are first lysed, mRNA is then isolated using magnetic beads, and in turn reverse transcribed into cDNA. Following cDNA synthesis, all samples are pooled, libraries are made, and the samples are sequenced. The protocol has been validated on 17 organisms, including human, mouse, zebrafish, and arabidopsis. Additionally, prime-seq is sensitive and works with low inputs, and the affordability of the method allows one to increase sample size to gain more biological insight. To verify prime-seq’s performance, we first compared prime-seq to TruSeq using the publicly available MAQC-III Study data. We then showed robust detection of marker genes in NPC differentiation and high throughput analysis of AML-PDX patient samples without compromising the archived samples.

## Results

### Development of the prime-seq protocol

The prime-seq protocol is based on the scRNA-seq method SCRB-seq [44] and our optimized derivative mcSCRB-seq [11]. It uses the principles of poly(A) priming, template switching, early barcoding, and UMIs to generate 3’ tagged RNA-seq libraries (Figure 1 and Figure S1). Compared to previous versions as described e.g. in [32], we have optimized the workflow, switched from a Nextera library preparation protocol to an adjusted version of NEBNext Ultra II FS, and made the sequencing layout analogous to 10X Chromium v3 gene expression libraries to facilitate pooling of libraries on Illumina flow cells, which is of great practical importance [16]. A detailed step-by-step protocol of prime-seq, including all materials and expected results, is available on protocols.io (https://dx.doi.org/10.17504/protocols.io.s9veh66). We have so far used this and previous versions of the protocol in 22 publications [9,23–43] and have generated just within the last year over 24 billion reads from >4,800 RNA-seq libraries in 97 projects from vertebrates (mainly mouse and human), plants, and fungi (Table S1 and Figure 2A). From these experiences, we find that the protocol works robustly and detects per sample on average >20,000 genes with 6.7 million reads of which 90.0% map to the genome and 71.6% map to exons and introns (Table S1). Notably, a large fraction (21%) of all UMIs map to introns with considerable variation among samples (Figure 2A).

**Figure 2.**
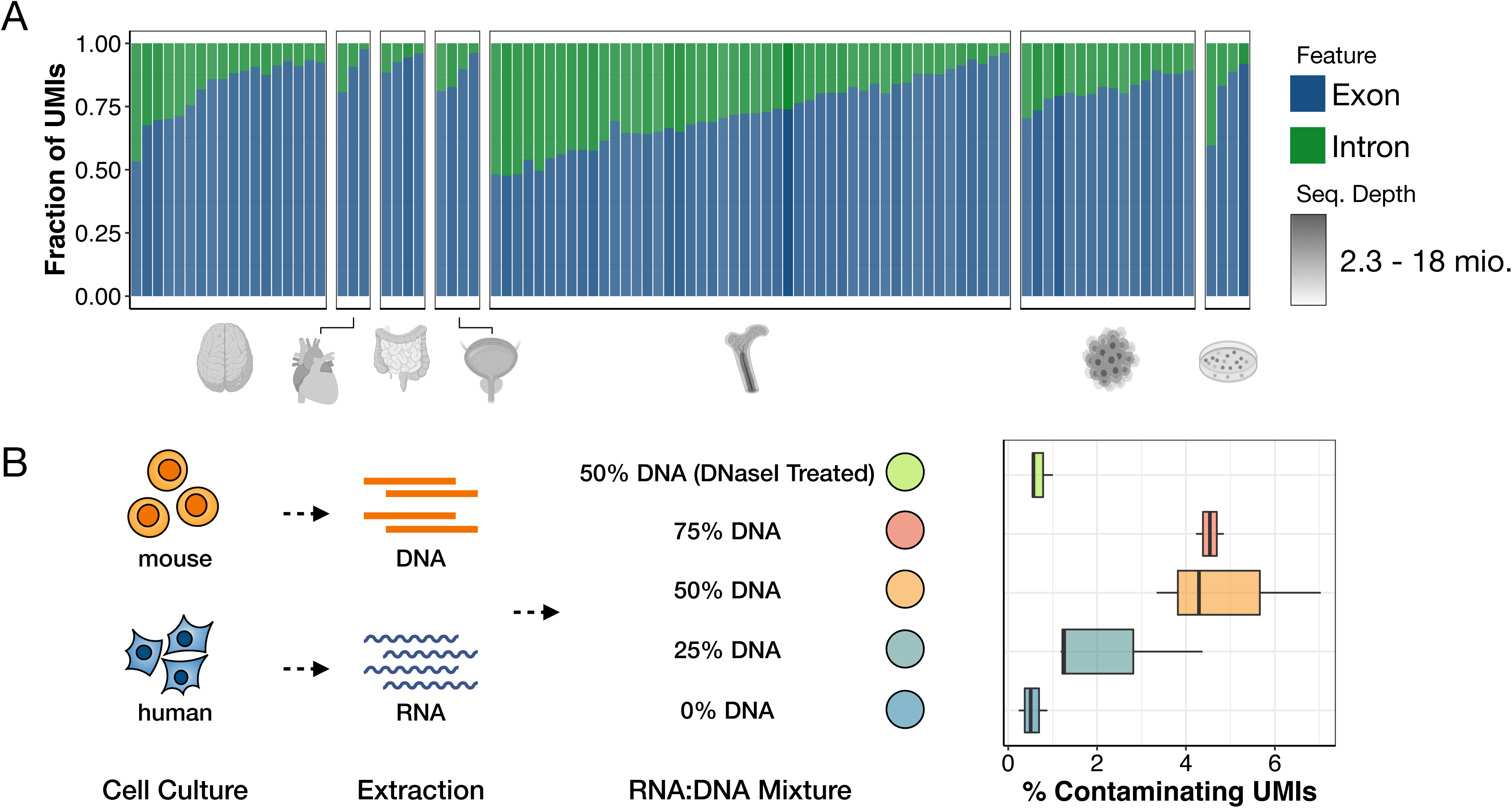
Intronic reads account for a variable but substantial fraction of UMIsand stem from RNA. (A) Fraction of exonic and intronic UMIs from 97 primate and mouse experiments using various tissues (neural, cardiopulmonary, digestive, urinary, immune, cancer, induced pluripotent stem cells). Sequencing depth is indicated by shading of the individual bars. We observe an average of 21% intronic UMIs, with some level of tissue-specific deviations as e.g. immune cells generally have higher fractions of intronic reads. (B) To determine if intronic reads stem from genomic DNA or mRNA, we extracted DNA from mouse embryonic stem cells (mESCs) and RNA from human induced pluripotent stem cells (hiPSCs) and then pooled the two in various ratios (75, 50, 25, and 0% gDNA) and counted the percentage of genomic (=mouse-mapped) UMIs. This indicates that DNAse I treatment in prime-seq is complete and that observed intronic reads are derived from RNA.

About 8,000 genes are detected only by exonic reads, ∼ 8,000 by exonic and intronic reads, and ∼ 4,000 by intronic reads only (Figure 2B, Table S1). Intronic reads correlate well with exonic reads of the same gene in scRNA-seq [45] and bulk RNA-seq data sets [46] and intronic reads are also used to infer expression dynamics in scRNA-seq data [47]. Hence, intronic reads can in principle be informative for quantifying gene expression. However, it is an uncommon practice to use them. This might be due to concerns that intronic reads could at least partially be derived from genomic DNA as MMLV-type reverse transcriptases could prime DNA that escaped a DNase I digest. Therefore, we investigated the origin of the intronic reads in prime-seq.

### Intronic reads are derived from RNA

First, we measured the amount of DNA yield generated from genomic DNA (gDNA). We lysed varying numbers of cultured human embryonic kidney 293T (HEK293T) cells and treated the samples with DNase I, RNase A, or neither prior to cDNA generation using the prime-seq protocol (up to and including the pre-amplification step). Per 1,000 HEK cells, this resulted in ∼5 ng of “cDNA” generated from gDNA in addition to the 12-32 ng of cDNA generated from RNA. (Figure S3A). To test the efficiency of DNase I digestion and quantify the actual number of reads generated from gDNA, we mixed mouse DNA and human RNA in different ratios (Figure 2B). Prime-seq libraries were generated and sequenced from untreated and DNAse I treated samples and reads were mapped to the mouse and human genome (Figure 2B). In the sample that did not contain any mouse DNA, ∼0.5% of all exonic and intronic UMIs mapped to the mouse genome, which represents the background level due to mismapping. Of the human mapped reads in this sample, ∼70% mapped to exons or introns and 10% to intergenic regions. (Figure S3B). Importantly, the DNAse I treated sample had the same distribution of mapped UMIs (0.7% mapped to mouse), strongly suggesting that the DNAse I digest is nearly complete and that essentially all reads in the DNAse I treated sample are derived from RNA.

As expected, with increasing amounts of mouse DNA the proportion of mouse mapped UMIs increased (Figure 2B), but even with 75% of the sample being mouse DNA, only 4.5% of the UMIs map to the mouse genome, suggesting that also for gDNA containing samples the impact of genomic reads on expression levels is likely small. Notably, with increasing amounts of gDNA, the fraction of unmapped reads also increased (Figure S3B), suggesting that the presence of gDNA does decrease the quality of RNA-seq libraries and does influence which molecules are generated during cDNA generation. In summary, these results indicate that essentially all reads in prime-seq libraries are derived from RNA when samples are DNAse I treated and hence that intronic reads can be used to quantify expression levels.

### prime-seq performs as well as TruSeq

Next, we quantitatively compared the performance of prime-seq to a standard bulk RNA-seq method with respect to library complexity, accuracy, and statistical power. A gold standard RNA-seq data set was generated in the third phase of the Microarray Quality Control (MAQC-III) study [48], consisting of deeply sequenced TruSeq RNA-seq libraries generated from five replicates of Universal Human Reference RNA (UHRR) and External RNA Controls Consortium (ERCC) spike-ins. As Illumina’s TruSeq protocol can be considered a standard bulk RNA-seq method and as the reference RNAs (UHRR and ERCCs) are commercially available, this is an ideal data set to benchmark our method. As in the MAQC-III design, we mixed UHRR and ERCCs (Figure S4A) in the same ratio but at a 1,000-fold lower input and generated eight prime-seq libraries, which were sequenced to a depth of at least 30 million reads. We processed and downsampled both data using the zUMIs pipeline [45] and compared the two methods with respect to their library complexity (number and expression levels of detected genes), accuracy (correlation of estimated expression level and actual number of spiked-in ERCCs), and statistical power (true positive and false positive rates in data simulated based on the mean-variance distribution of technical replicates of each method).

We found that prime-seq has a slightly lower fraction of exonic and intronic reads that can be used to quantify gene expression (78% vs. 85%; Figure 3A, Figure S5A). But despite the slightly lower number of reads that can be used, prime-seq does detect at least as many genes as TruSeq (Figure 3B). Both methods also show a similar distribution of gene expression levels (Figure 3D), indicating that the complexity of generated libraries is generally very similar.

**Figure 3.**
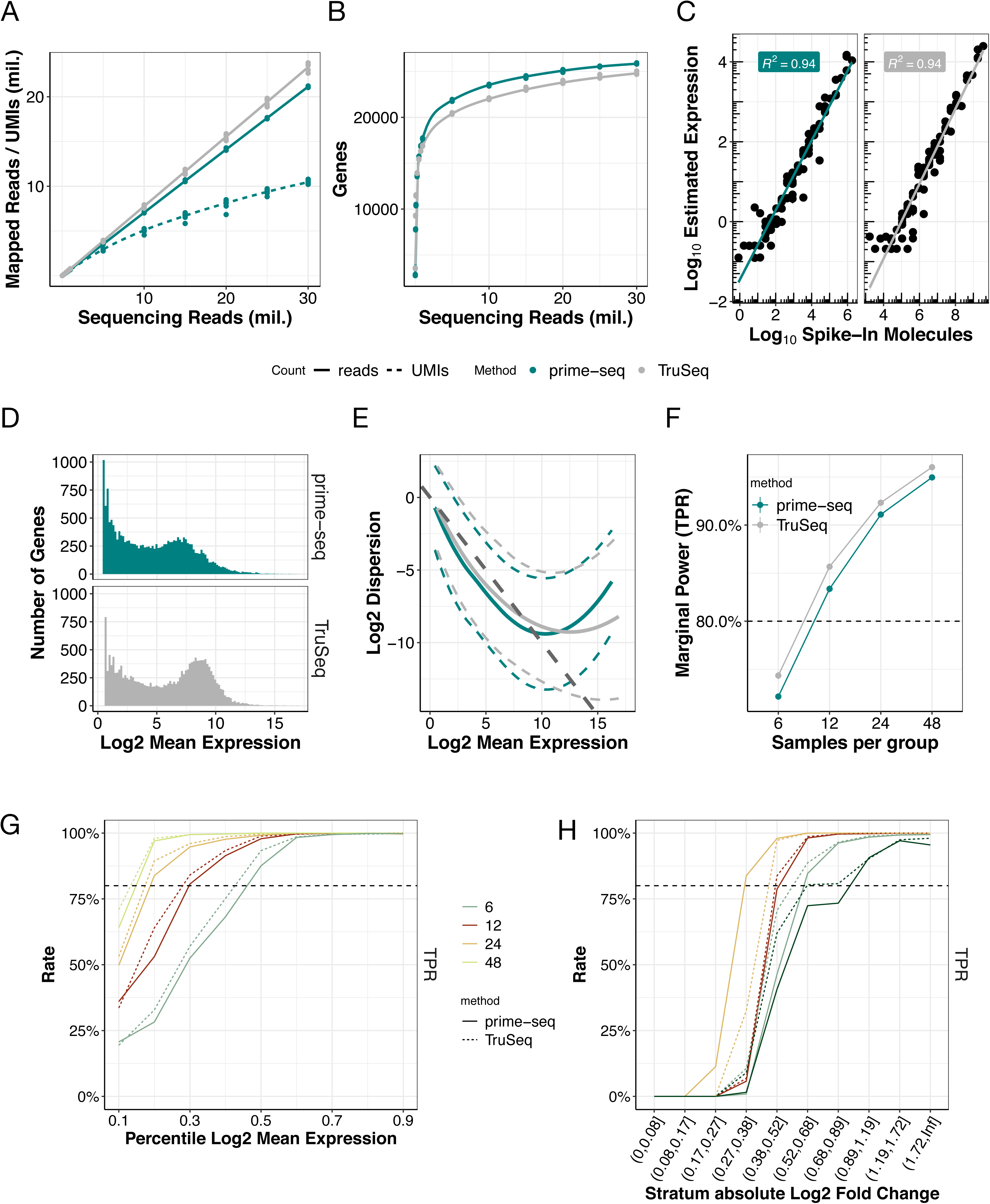
prime-seq has similar sensitivity and power compared to TruSeq (MAQC-III data). (A) Mapped reads, UMIs (dashed line, only prime-seq), and (B) detected genes at varying sequencing depths between TruSeq data from the MAQC-III Study and matched prime-seq data, shows prime-seq and TruSeq are similarly sensitive (filtering parameters: detected UMI ≥ 1, detected gene present in at least 25% of samples and is protein coding). (C) Accuracy, measured by spike-in molecules, is similarly high in both methods (R^2^=0.94). (D) The distribution of genes across mean expression is similar for both methods, as well as the (E) dispersion, which follows a poisson distribution (dark grey dashed line) for lower expressed genes and then increases as technical variation increases for highly expressed genes. The local polynomial regression fit between mean and dispersion estimates per method is shown in solid lines with 95% variability band per gene shown in dashed lines. (F) Power analysis at a sequencing depth of 10 million reads shows almost identical power between prime-seq and TruSeq, and a similar increase at varying sample size for (G) mean expression and (H) absolute log2 fold change. Data filtering parameters: detected UMI ≥ 1, detected gene present in at least 25% of samples.

The accuracy of a method, i.e. how well estimated expression levels reflect actual concentrations of mRNAs, is relevant when expression levels are compared among genes. Here, TruSeq and prime-seq show the same correlation (Pearson’s R^2^ = 0.94) between observed expression levels and the known concentration of ERCC spike-ins, indicating that their accuracy is very similar (Figure 3C).

However, for most RNA-seq experiments, a comparison among samples - e.g. to detect differentially expressed genes - is more relevant. Therefore, it matters how well genes are measured by a particular method, i.e. how much technical variation a method generates across genes. As we have 8 and 5 technical replicates of the same RNA for prime-seq and TruSeq, respectively, we can estimate for each method the mean and variance per gene. Note that UMIs are only available for prime-seq and hence only prime-seq can profit from removing technical variance by removing PCR duplicates (Figure 3A). The empirical distribution shows the characteristic dependency of RNA-seq data on sampling (Poisson expectation) at low expression levels and an increasing influence of the additional technical variation at higher expression levels (Figure 3E). Prime-seq shows a slightly lower variance for medium expression levels where most genes are expressed and a higher one for a handful of genes with very high expression levels (Figure 3E). To quantify to what extent these differences in the mean-variance distribution actually matter, we used power simulations as implemented in powsimR [49]. We simulated that 10% of genes sampled from the estimated mean-variance relation of each method are differentially expressed between two groups of samples. The fold changes of these genes were drawn from a distribution similar to those we observed in actual data between two cell types (iPSCs and NPCs) or two types of acute myeloid leukemia (AML) (see below and Figure S5B). The comparison between this ground truth and the identified differentially expressed genes in a simulation allows us to estimate the true positive rate (TPR) and the false discovery rate (FDR) for a particular parameter setting. We stratified TPR and FDR across the number of replicates (Figure 3F), the expression levels (Figure 3G), and the fold changes (Figure 3H) to illustrate the strong dependence of power on these parameters. At a given FDR level, a more powerful method reaches a TPR of 80% with fewer replicates, at a lower expression level, and/or for a lower fold change. We find that the power of the two methods is almost identical as FDR and TPR are very similar across conditions for both methods. The false discovery rates (FDR) are - as expected - generally below 5% for 12, 24, or 48 replicates per condition (Figure S5C) and the (marginal) TPR across all expression levels and fold changes is 80% for both methods at ∼12 replicates per condition (Figure 3F). The power increases for both methods in a similar manner with increasing expression levels (Figure 3G) and increasing fold changes (Figure 3H). This is also the case when using only exonic reads for the power analysis (Figure S5C and S5F-G). In summary, prime-seq and TruSeq perform very similarly in estimating gene expression levels with respect to library complexity, accuracy, and statistical power.

### Bead-based RNA extraction increases cost efficiency and throughput

As library costs and sequencing costs drop, standard RNA isolation becomes a considerable factor for the cost efficiency of RNA-seq methods. RNA isolation using magnetic beads is an attractive alternative [50] and we have used it successfully in combination with our protocol before [11]. To investigate the effects of RNA extraction more systematically, we compared prime-seq libraries generated from RNA extracted via silica columns and via magnetic beads. Libraries from cultured HEK293T cells, human peripheral blood mononuclear cells (PBMC), and mouse brain tissue showed a similar distribution of mapped reads, albeit with a slightly higher fraction of intronic reads in magnetic bead libraries (Figure 4A and S6) and considerable differences in expression levels (Figure 4B and S7).

**Figure 4.**
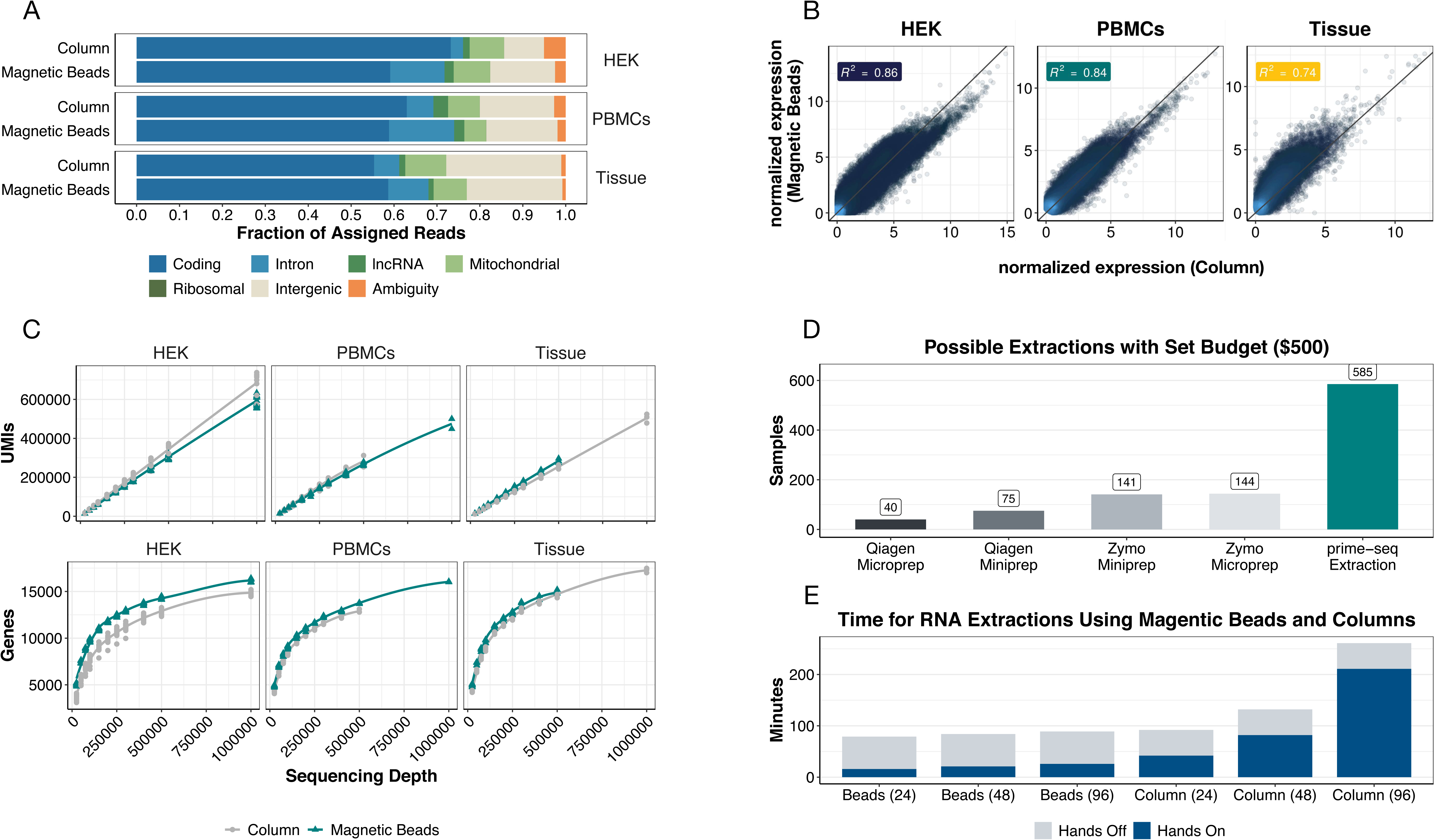
RNA extraction with beads, rather than columns, provides similar sequencing data while increasing throughput capabilities. (A) Feature distributions of RNA isolated with a column-based kit and magnetic beads show that both RNA extraction protocols produce similar amounts of useable reads from cultured human embryonic kidney 293T (HEK293T) cells, peripheral blood mononuclear cells (PBMC), and harvested mouse brain tissue. (B) Gene expression between both bead and column extraction are also similar in all three tested inputs (R^2^ = 0.86 HEK, 0.84 PBMCs, and 0.74 tissue). (C) Detected UMIs and detected genes for column and magnetic beads in HEK293T, PBMCs, and tissue are almost identical, with slightly more detected genes in the bead condition (filtering parameters: detected UMI ≥ 1, detected gene present in at least 25% of samples and is protein coding). Comparison of costs (D) and time (E) required for different RNA extractions.

To further explore these differences, we tested the influence of the Proteinase K digestion and its associated heat incubation (50°C for 15 minutes and 75°C for 10 minutes), which is part of the bead based RNA isolation protocol. We prepared prime-seq libraries using HEK293T RNA extracted via silica-columns (“Column”), magnetic beads with Proteinase K digestion (“Magnetic Beads”), magnetic beads without Proteinase K digestion (“No Incubation”), and magnetic beads with the same incubations but without the addition of the enzyme (“Incubation”). Interestingly, the shift to higher intronic fractions and the expression profile similarity is mainly due to the heat incubation, rather than the enzymatic digestion by Proteinase K (Figure S6A and B).

Hence, bead-based extraction does create a different expression profile than column based extraction, especially due to the often necessary Proteinase K incubation step. This confirms the general influence of RNA extraction protocols on gene expression profiles [51]. Importantly, the complexity of the two types of libraries is similar, with a slightly higher number of genes detected in the bead-based isolation (Figure 4C, Figure S6C and S6D), potentially due to a preference for longer transcripts with lower GC contents (Figure S7C).

So while bead-based RNA isolation and column-based RNA isolation create different but similarly complex expression profiles, bead-based RNA isolation has the advantage of being much more cost-efficient. At least four times more RNA samples can be processed for the same budget (Figure 4D, Table S2). In addition, RNA isolation using magnetic beads is twice as fast and without robotics more amenable to high throughput experiments (Table S3). Thus, we show that bead-based RNA isolation can make prime-seq considerably more cost-efficient without compromising library quality.

### prime-seq is sensitive and works well with 1,000 cells

As prime-seq was developed from a scRNA-seq method [44], it is very sensitive, i.e. it generates complex libraries from one or very few cells. This makes it useful when input material is limited, e.g. when working with rare cell types isolated by FACS or when working with patient material. To validate a range of input amounts, we generated RNA-seq libraries from 1,000 (low input, ∼10-20 ng total RNA) and 10,000 (high input, ∼100-200 ng) HEK293T cells. The complexity of the two types of libraries was very similar, with only a 2% decrease in the fraction of exonic and intronic reads and a 7.7% and 1.9% reduction in the number of UMIs and detected genes at the same sequencing depth (Figure S8A). The expression profiles were almost as similar between the two input conditions as within the input conditions (median r within = 0.94, median r between = 0.93; Figure S8B), indicating that expression profiles from 1,000 and 10,000 cells are almost identical in prime-seq. Using a lower number of input cells is certainly possible and unproblematic as long as the number of cells is unbiased with respect to the variable of interest. Using higher amounts than 10,000 cells is certainly also possible, but it is noteworthy that we have observed a large fraction of intergenic reads in highly concentrated samples, potentially due to incomplete DNase I digestion (data not shown). In summary, we validate that an input amount of at least 1,000 cells does not compromise the complexity of prime-seq libraries and hence that prime-seq is a very sensitive RNA-seq protocol.

### Two exemplary applications of prime-seq

To exemplify the advantages with respect to sensitivity and throughput in an actual setting, we used prime-seq to profile cryopreserved human acute myeloid leukemia (AML) cells from patient-derived xenograft (PDX) models [23,52]. These consisted of different donors and AML subtypes and were stored in freezing medium at -80°C for up to 3.5 years (Figure 5A). Due to the sensitivity of prime-seq, we could use a minimal fraction of the sample without thawing it by taking a 1 mm biopsy punch from the vial of cryopreserved cells and putting it directly into the lysis buffer. This allowed sampling of precious samples without compromising their amount or quality and resulted in 94 high quality expression profiles that clustered mainly by AML subtype (Figure 5B) as expected [53].

**Figure 5.**
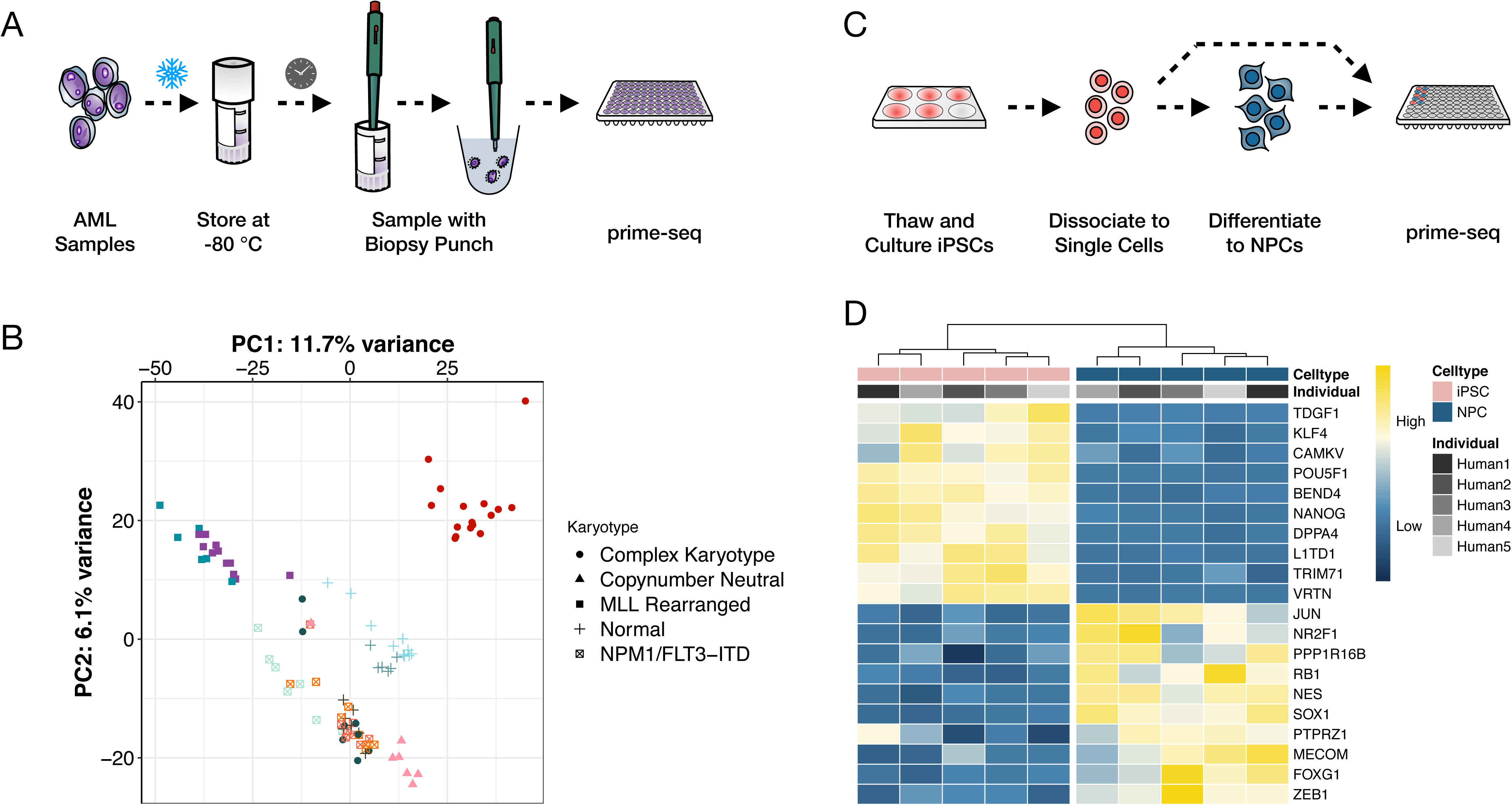
Two exemplary applications of prime-seq. (A) Experimental design for an acute myeloid leukemia (AML) study, where a biopsy punch was used to collect a small fraction of a frozen Patient-derived xenograft (PDX)-AML sample. (B) Prime-seq libraries were generated from 94 PDX samples, derived from 11 different AML-PDX lines (colour-coded) from 5 different AML subtypes (symbol-coded) and cluster primarily by AML subtype. (C) Experimental design for studying the differentiation from five human induced pluripotent stem cell lines (iPSCs) to neural progenitor cells (NPC). (D) Expression levels from 20 a priori known marker genes cluster iPSCs and NPCs as expected.

To further exemplify the performance of prime-seq, we investigated its ability to detect known differences in a well established differentiation system [54]. We differentiated five human induced pluripotent stem cell (iPSCs) lines [36] to neural progenitor cells (NPCs) and generated expression profiles using prime-seq (Figure 5C). In a hierarchical clustering of well known marker genes [55], the iPSCs and NPCs formed two distinct groups and the expression patterns were in agreement with their cellular identity. For example the iPSC markers POU5F1, NANOG and KLF4 showed an increased expression in the iPSCs and NES, SOX1, and FOXG1 in NPCs (Figure 5D).

### prime-seq is cost-efficient

We have shown above that the power, accuracy and library complexity is similar between prime-seq and TruSeq. The performance and robustness of the prime-seq protocol has been demonstrated by the two examples above as well as its many applications using this or previous versions of the protocol [9,23–35,42,43,56,57]. In summary, one could argue that prime-seq performs as well as TruSeq for quantifying gene expression levels. Other methods that generate tagged cDNA libraries using early barcoding have also been developed [16,22,58–61]. This includes BRB-seq that uses poly(A) priming and DNA-Pol I for second strand synthesis and also performs similarly to TruSeq [22]. Decode-seq also uses poly(A) priming and template switching like prime-seq, but adds sample-specific barcodes and UMIs at the 5’ end [16]. In a direct comparison, Decode-seq performed slightly better than BRB-seq and due to a more flexible sequencing layout [16]. While slight differences in power, accuracy, and/or library complexity might exist among these protocols, cross-laboratory benchmarking on exactly the same samples as recently done e.g. for scRNA-seq methods [5] or small RNA-seq methods [62] are probably needed to quantify such differences reliably. For now, it is probably fair to say that RNA-seq methods like BRB-seq, prime-seq, TruSeq, SmartSeq, or Decode-seq all perform fairly equal with respect to quantifying gene expression levels. Hence, at a fixed budget the cost per sample will determine to a large extent how many samples can be analyzed and hence how much biological insight can be gained.

To this end, we calculated the required reagent costs to generate a library from isolated RNA in a batch of 96 samples for the different commercial methods as well as for prime-seq, Decode-seq, and BRB-seq (Table S4). With $2.53 per sample prime-seq is the most cost-efficient method, followed by BRB-seq ($4.05) and Decode-seq ($6.58). Commercial methods range from $60 (NEBNext) to $164 (SMARTer Stranded). This is illustrated by the number of libraries that can be generated by a fixed budget of $500 (Figure 6A). Note that these costs include for all methods $1.39 per sample for two Bioanalyzer (Agilent) Chips (Table S4) and do not consider the additional cost reduction that is associated with the direct bead-based RNA extraction of prime-seq (see above). The drastic advantage of prime-seq, Decode-seq, and BRB-seq also becomes apparent when power is plotted as a function of costs with and without sequencing (10 million reads per sample) (Figure 6B, Figure S9A). For example, to reach an 80% TPR at a desired FDR of 5%, one needs to spend $715 including sequencing costs for prime-seq, $795 when using Decode-seq, $1,625 when using Illumina Stranded, and $3,485 when using TruSeq (Figure S9B).

**Figure 6.**
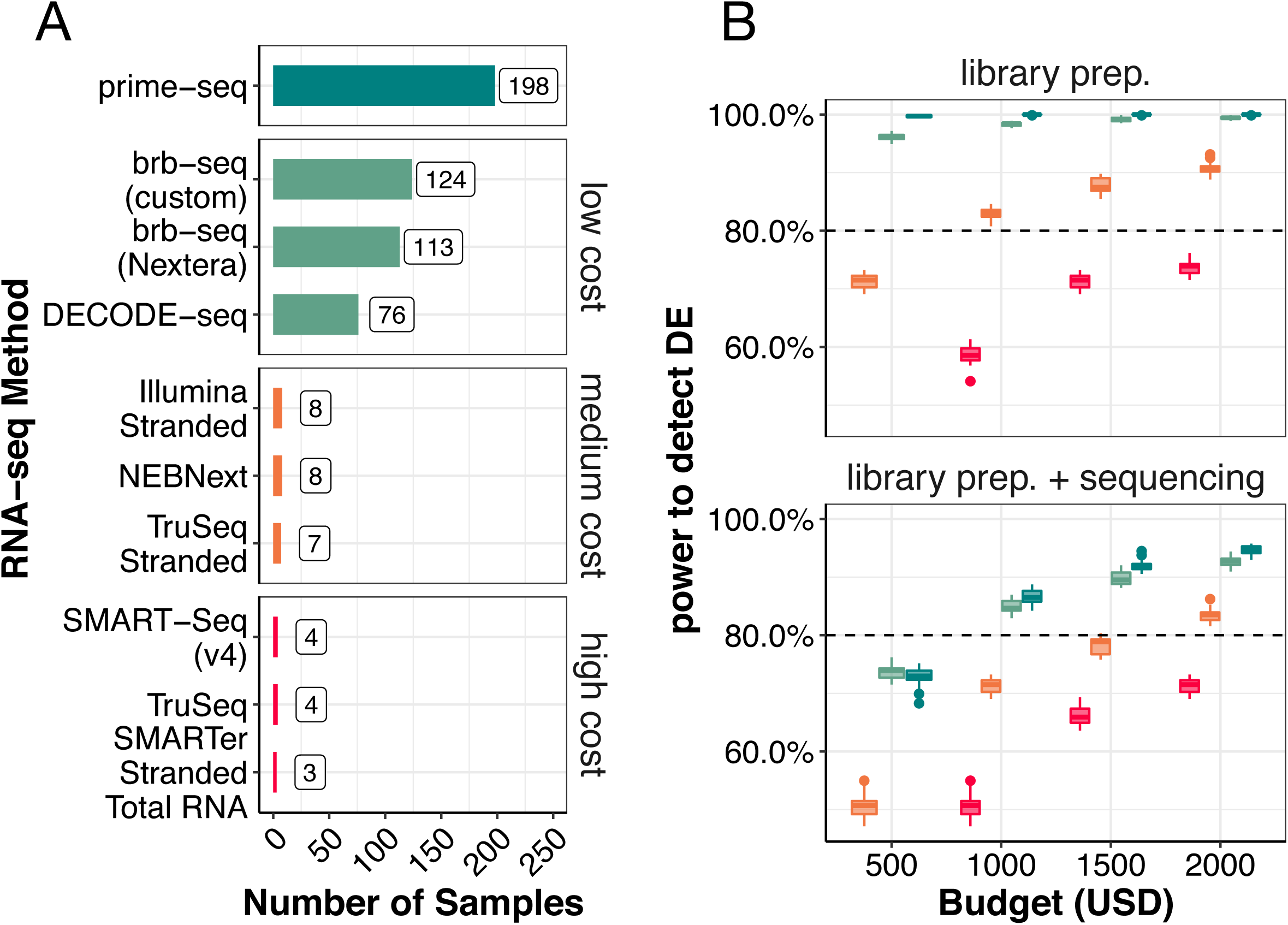
Prime-seq is very cost-efficient. (A) With a set budget of $500, prime-seq allows one to process 198 samples, which is 1.6 times more samples than the next cost-efficient method. (B) The compared methods were grouped into low, middle, and high cost methods and the TruSeq MAQCII data was used as a basis for power analysis for all methods but prime-seq. The increase in sample size due to cost efficiency directly impacts the power to detect differentially expressed genes, as evident by the increased performance of prime-seq and other low cost methods (BRB-seq and Decode-seq), even when sequencing costs are included in the comparison (sequencing depth of 10 mio. reads at a cost of $3.40 per 1 mio. reads).

Cost-efficiency with respect to time can also matter and we calculated hands-on and hands-off time for the different methods (Table S5). Hands-on times vary from 30-35 minutes for the non-commercial, early barcoding methods to 52-191 minutes for commercial methods. However, as all methods require essentially a full day of lab work, we consider the differences in required times not as decisive, at least not in a research lab setting where RNA-seq is not done on a daily or weekly basis. In summary, we find that prime-seq is the most cost-efficient bulk RNA-seq method currently available.

## Discussion

In this paper we present and validate prime-seq, a bulk RNA-seq protocol, and show that it is as powerful and accurate as TruSeq in quantifying gene expression levels, but more sensitive and much more cost-efficient. We validate the DNAse I treatment and determine that intronic reads are derived from RNA and can be used in downstream analysis. We also validate input ranges and the direct lysis and bead-based RNA purification of tissue and cell culture samples. Finally, we exemplify the use of prime-seq by profiling AML samples and NPC differentiation and show that prime-seq is currently the most cost-efficient bulk RNA-seq method. In the following, we focus our discussion on advantages and drawbacks of prime-seq in comparison to other RNA-seq protocols. To this end, we distinguish protocols like TruSeq, Smart-seq, or NEBNext that individually process RNA samples and generate full-length cDNA profiles (“full-length protocols”) from protocols like prime-seq, Decode-seq, or BRB-seq that use early barcoding and generate 5’ or 3’ tagged cDNA libraries (“tag protocols”).

### Complexity, power and accuracy are similar among most bulk RNA-seq protocols

Initially, early barcoding 3’ tagged protocols generated slightly less complex libraries (i.e. detected fewer genes for the same number of reads), especially due to a considerable fraction of unmapped reads [22,63]. These reads are probably caused by PCR artifacts during cDNA generation and amplification. Protocol optimizations as shown for BRB-seq [22], Decode-seq [16] and here for prime-seq have reduced these artifacts and hence have improved library complexity to the level of standard full-length protocols. For prime-seq we have shown quantitatively that its complexity, accuracy, and power is very similar to that of TruSeq. More comprehensive studies, ideally across laboratories [5,48], would be needed to quantitatively compare protocols, also with respect to their robustness across laboratories and conditions and their biases for individual transcripts. For the context and methods discussed here, we would argue that there are no decisive differences in power, accuracy, and complexity among tag protocols and full-length protocols at least when performed under validated and optimized conditions.

### Cost-efficiency makes tag-protocols preferable when quantifying gene expression levels

As shown above (Figure 6) and as argued before [16,22,63], the main advantage of tag protocols is their cost-efficiency. Their most obvious drawback is that they cannot quantify expression levels of different isoforms. Smart-seq2 [64] and Smart-seq3 [10] are relatively cost-efficient full length protocols that were developed for scRNA-seq. However, they have not been validated and optimized for bulk RNA-seq and would still be considerably more expensive than most tag protocols. Furthermore, as reconstructing transcripts from short read data is difficult and requires deep sequencing, isoform detection and quantification is now probably more efficiently done by using long-read technologies [1]. However, from our experience, most RNA-seq projects quantify expression at the gene level not at the transcript level. This is probably because most projects use RNA-seq to identify affected biological processes or pathways by a factor of interest. As different genes are associated with different biological processes, but different isoforms are only very rarely associated with different biological processes, most projects do not profit much from quantifying isoforms. Hence, we would argue that quantifying expression levels of genes is the better option, as long as isoform quantification is not of explicit relevance for a project.

Another limitation is that all tag-protocols use poly(A) priming and hence do not capture mRNA from bacteria, organelles, or other non-polyadenylated transcripts. For full-length protocols like TruSeq, cDNA generation by random priming after rRNA depletion can be done. Another possibility is poly(A) tailing after rRNA depletion [65], but to our knowledge, this has not been adopted to tag-based protocols yet. How to efficiently combine profiling of polyadenylated, non-polyadenylated, and small RNA is certainly worth further investigating. However, it is also true that for eukaryotic cells, quantification of mRNAs contains most of the information. Hence, similar to the quantification of isoforms, we would argue that quantifying expression levels of genes by polyadenylated transcript is often sufficient, as long as non-polyadenylated transcripts are not explicitly relevant.

Finally, while early barcoding and pooling enable the cost-efficiency of tag protocols, this necessitates calibrating input amounts. Input calibration is easy when starting with extracted RNA or when it is possible to count cells prior to direct lysis. When counting cells is not possible, we have also developed a protocol adaptation of prime-seq that allows for RNA quantification and normalization after bead-based RNA isolation and prior to reverse transcription (https://dx.doi.org/10.17504/protocols.io.s9veh66). Early barcoding and pooling also entails the danger of barcode swapping, i.e. the formation of chimeric molecules during PCR, resulting in a contamination of a cell’s expression profile with transcripts from another cell. This is especially an issue for scRNA-seq [66] as the number of PCR cycles and on the polymerase likely play a role [67]. To verify that this is not an issue in prime-seq, we pooled human and mouse samples at each possible point in the protocol; we detected low rates of cross-contamination when samples were pooled as RNA (0.59%), cDNA (0.76%), or libraries (0.83%).

In summary, when quantification of isoforms and/or non-polyadenylated RNA is not necessary, a technically validated tag protocol has no drawbacks. Protocols that use poly(A) priming and template switching also have the advantage that they are very sensitive and for prime-seq we have validated that it still works optimally also with 1,000 cells (∼10-20ng total RNA) as input. However, the decisive advantage of tag protocols is their drastically higher cost-efficiency (Figure 6), as this leads to drastically higher power and much more flexibility in the experimental design for a given budget. As repeated by biostatisticians over the decades, a good experimental design and a sufficient number of replicates is the most decisive factor for expression profiling. It is sobering how enduring the n=3 tradition is, as is nicely shown in [16], although it is known that it is better to distribute the same number of reads across more biological replicates [17]. Cost-efficient tag protocols will hopefully make such experimental designs more common. While library costs are less notable for sequencing depths of 10M reads or more (Figure 6B), they may enable RNA-seq experiments that can be done with shallow sequencing, something which is less obvious and might be overlooked. Replacing qPCR has been advocated as one example by the authors of BRB-seq[22]. But also other applications, like characterizing cell type composition [36], quality control of libraries, or optimizing experimental procedures can profit considerably from low library costs.

In summary, tag protocols allow flexible designs of RNA-seq experiments that should be helpful for many biological questions and have a vast potential when readily accessible for many labs.

### Validation, documentation, and cost-efficiency make prime-seq a good option for setting up a tag protocol

We have argued above that adding a tag protocol to the standard method repertoire of a molecular biology lab is advantageous due to its cost-efficiency. As the different tag protocols discussed here perform fairly similar with respect to complexity, power, accuracy, sensitivity, and cost-efficiency, essentially any of them would suffice. If one has a validated, robust protocol running in a lab or core facility, it is probably not worth switching. That said, our results might still help to better validate existing protocols, integrate direct lysis, and make use of intronic reads. If one does not have a tag protocol running, we would argue that our results provide helpful information to decide on a protocol, and that prime-seq would be a good option for several reasons as laid out in the following.

A main difference among tag protocols is whether they tag the 5’ end, like Decode-seq, or tag the 3’ end like BRB-seq or prime-seq. 5’ tagging has some obvious advantages (see also [16]), including the possibility to read both ends of the cDNA as one cannot read through the poly(A) tail. Using the sequence information from the 5’ end is also important to distinguish alleles of B-cell receptors and T-cell receptors [68]. In scRNA-seq, both 5’ and 3’ tag protocols have been successfully used, but 3’ tagging is currently the standard. The reason for this is not obvious, but it might be that the incorporation of the barcode and the UMI is more difficult to optimize [10]. Additionally, the higher level of alternative splicing at the 5’ end could make gene-level quantification more difficult. More dedicated comparisons would be needed to further investigate these factors. Currently, 3’ tag protocols are more established and when using a suitable sequencing design, poly(A) priming does not compromise sequencing quality as validated by us and the widespread use of Chromium 10x v3 chemistry scRNA-seq libraries that have the same layout as prime-seq.

As shown above, prime-seq is among all protocols the most cost-efficient when starting from purified RNA. It is also currently the only protocol for which a direct lysis is validated, which further increases cost-efficiency of library production. This is especially advantageous when processing many samples, shallow sequencing is sufficient, and/or as sequencing costs continue to drop.

Finally, we think that prime-seq is the easiest tag protocol to set up. While many such protocols have been published and all have argued that their method would be useful, few have actually become widely implemented. The reasons are in all likelihood complex, but we think that prime-seq has the lowest barriers to be set up by an individual lab or a core facility for three reasons: First, to our knowledge it is the most validated non-commercial bulk RNA-seq protocol, based on the experiments presented here as well as our >5 years of experience in running various versions of the protocol with over 6,000 samples across 17 species resulting in over 20 publications to date. It is the only protocol for which direct lysis and sensitivity are quantitatively validated. Also, it is well validated in combination with zUMIs, the computational pipeline that was developed and is maintained by our group [45]. Second, it is not only cost-efficient per sample, but it also has low setup costs. It requires no specialized equipment and only the barcoded primers as an initial investment of ∼$2,000 for 96 primers, which will be sufficient for processing more than 240 thousand samples. Finally, prime-seq is well documented not only by this manuscript, but also by a step-by-step protocol, including all materials, expected results, and alternative versions depending on the type and amounts of input material (https://dx.doi.org/10.17504/protocols.io.s9veh66). Hence, we think that prime-seq is not only a very useful protocol in principle, but also in practice.

## Conclusion

The multi-dimensional phenotype of gene expression is highly informative for many biological and medical questions. As sequencing costs dropped, RNA-seq became a standard tool in investigating these questions. We argue that the decisive next step is to use the possibilities of lowered library costs by tag protocols to leverage even more of this potential. We show that prime-seq is currently the best option when establishing such a protocol as it performs as well as other established RNA-seq protocols with respect to its accuracy, power, and library complexity. Additionally, it is very sensitive, is well documented, and is the most cost-efficient bulk RNA-seq protocol currently available to set up and to run.

## Methods

A step-by-step protocol of prime-seq, including all materials and expected results, is available on protocols.io (https://dx.doi.org/10.17504/protocols.io.s9veh66). Below, we briefly outline the prime-seq protocol, as well as describe any experiment-specific methods and modifications that were made to prime-seq during testing and optimization.

### prime-seq

Cell lysates, generally containing around 1,000-10,000 cells, were treated with 20 µg of Proteinase K (Thermo Fisher, #AM2546) and 1µL 25 mM EDTA (Thermo Fisher, EN0525) at 50°C for 15 minutes with a heat inactivation step at 75°C for 10 minutes. The samples were then cleaned using cleanup beads, a custom made mixture containing SpeedBeads (GE65152105050250, Sigma-Aldrich), at a 1:2 ratio of lysate to beads. DNA was digested on-beads using 1 unit of DNase I (Thermo Fisher, EN0525) at 20°C for 10 minutes with a heat inactivation step at 65°C for 5 minutes.

The samples were then cleaned and the RNA was eluted with the 10 µL reverse transcription mix, consisting of 30 units Maxima H- enzyme (Thermo Fisher, EP0753), 1x Maxima H- Buffer (Thermo Fisher), 1 mM each dNTPs (Thermo Fisher), 1 µM template-switching oligo (IDT), 1 µM barcoded oligo(dT) primers (IDT). The reaction was incubated at 42°C for 90 minutes.

Following cDNA synthesis, the samples were pooled, cleaned, and concentrated with cleanup beads at a 1:1 ratio and eluted in 17 µL of ddH2O. Residual primers were digested using Exonuclease I (Thermo Fisher, EN0581) at 37 °C for 20 minutes followed by a heat inactivation step at 80 °C for 10 minutes. The samples were cleaned once more using cleanup beads at a 1:1 ratio, and eluted in 20 µL of ddH2O.

Second strand synthesis and pre-amplification were performed in a 50 µL reaction, consisting of 1x KAPA HiFi Ready Mix (Roche, 7958935001) and 0.6 µM SingV6 primer (IDT), with the following PCR setup: initial denaturation at 98 °C for 3 minutes, denaturation at 98 °C for 15 seconds, annealing at 65 °C for 30 seconds, elongation at 68 °C for 4 minutes, and a final elongation at 72 °C for 10 minutes. Denaturation, annealing, and elongation were repeated for 5-15 cycles depending on the initial input.

The DNA was cleaned using cleanup beads at a ratio of 1:0.8 of DNA to beads and eluted with 10 µL of ddH2O. The quantity was assessed using a Quant-iT PicoGreen dsDNA assay kit (Thermo Fisher, P11496) and the quality was assessed using an Agilent 2100 Bioanalyzer with a High Sensitivity DNA analysis kit (Agilent, 5067-4626).

Libraries were prepared with the NEBNext Ultra II FS Library Preparation Kit (NEB, E6177S) according to manufacturer instructions in most steps, with the exception of adapter sequence and reaction volumes. Fragmentation was performed on 2.5 µL of cDNA (generally 2 - 20 ng) using Enzyme Mix and Reaction buffer in a 6 µL reaction. A custom prime-seq adapter (1.5 µM, IDT) was ligated using the Ligation Master Mix and Ligation Enhancer in a reaction volume of 12.7 µL. The samples were then double-size selected using SPRI-select Beads (Beckman Coulter, B23317), with a high cutoff of 0.5 and a low cutoff of 0.7. The samples were then amplified using Q5 Master Mix (NEB, M0544L), 1 µL i7 Index primer (Sigma-Aldrich), and 1 µL i5 Index primer (IDT) using the following setup: 98°C for 30 seconds; 10-12 cycles of 98°C for 10 seconds, 65°C for 1 minute 15 seconds, 65°C for 5 minutes; and 65°C for 4 minutes. Double-size selection was performed once more as before using SPRI-select Beads. The quantity and quality of the libraries were assessed as before.

### Nextera XT Library Prep

Prior to using the NEBNext Ultra II FS Library Kit, libraries were prepared using the Nextera XT Kit (Illumina, FC-131-1096). This included the RNA extraction experiments (Figure 4) as well as the AML experiment (Figure 5B). These libraries were prepared as previously described [11].

Briefly, three replicates of 0.8 ng of DNA were tagmented in 20 µL reactions. Following tagmentation, the libraries were amplified using 0.1 µM P5NextPT5 primer (IDT) and 0.1 µM i7 index primer (IDT) in a reaction volume of 50 µL. The index PCR was incubated as follows: gap fill at 72°C for 3 minutes, initial denaturation at 95 °C for 30 seconds, denaturation at 95 °C for 10 seconds, annealing at 62 °C for 30 seconds, elongation at 72 °C for 1 minute, and a final elongation at 72 °C for 5 minutes. Denaturation, annealing, and elongation were repeated for 13 cycles.

Size selection was performed using gel electrophoresis. Libraries were loaded onto a 2% Agarose E-Gel EX (Invitrogen, G401002) and were excised between 300 bp - 900 bp and cleaned using the Monarch DNA Gel Extraction Kit (NEB, T1020). The libraries were quantified and qualified using an Agilent 2100 Bioanalyzer with a High Sensitivity DNA analysis kit (Agilent, 5067-4626).

### Barcoded oligo(dT) primer design

In order to enable more robust demultiplexing and to ensure full compatibility of our sequencing layout with the Chromium 10x v3 chemistry, oligo(dT) primers were designed to include a 12 nt cell barcode and 16 nt UMI. Candidate cell barcodes were created in R using the DNABarcodes package [69] to generate barcodes with a length of 12 nucleotides and a minimum Hamming distance (HD) of 4, with filtering for self-complementarity, homo-triplets, and GC-balance enabled. Candidate barcodes were filtered further, resulting in a barcode pool with a minimal HD of 5 and a minimal Sequence-Levenshtein distance of 4 within the set. In order to balance nucleotide compositions among cell barcodes at each position BARCOSEL [70] was used to further reduce the candidate set down to the final 384 barcodes.

### Sequencing

Sequencing was performed on an Illumina HiSeq 1500 instrument for all libraries except for the IPSC/NPC experiment where a NextSeq 550 instrument was used. The following setup was used: Read 1: 28 bp, Index 1: 8 bp; Read 2: 50-56 bp.

### Pre-processing of RNA-seq Data

The raw data was quality checked using fastqc (version 0.11.8 [71]) and then trimmed of poly(A) tails using Cutadapt (version 1.12, https://doi.org/10.14806/ej.17.1.200). Following trimming, the zUMIs pipeline (version 2.9.4 ,[45]) was used to filter the data, with a Phred quality score threshold of 20 for 2 BC bases and 3 UMI bases. The filtered data was mapped to the human genome (GRCh38) with the Gencode annotation (v35) or the mouse genome (GRCm38) with the Gencode annotation (vM25) using STAR (version 2.7.3a,[72]) and the reads counted using RSubread (version 1.32.4,[73]).

### Sensitivity and Differential Gene Expression Analysis of RNA-seq Data

The count matrix generated by zUMIs was loaded into RStudio (version 1.3.1093 [74]) using R (version 4.0.3 [75]). bioMart (version 2.46.0 [76]), dplyr (version 1.0.2 [77]), and tidyr (version 1.1.2 [78]) were used for data processing and calculating descriptive statistics (i.e. detected genes, reads, and UMIs). DESeq2 (version 1.30.0 [79]) was used for differential gene expression analysis. ggplot2 (version 3.3.3 [80]), cowplot (version 1.1.1 [81]), ggbeeswarm (0.6.0 [82]), ggsignif (version 0.6.0 [83]), ggsci (version 2.9 [84]), ggrepel (version 0.9.0 [85]), EnhancedVolcano (1.8.0 [86]), ggpointdensity (version 0.1.0 [87]) and pheatmap (version 1.0.12 [88]) were used for data visualization.

### Power Analysis of RNA-seq Data

Power Simulations were performed following the workflow of the powsimR package (version 1.2.3 [49]). Briefly, RNAseq data per method was simulated based on parameters extracted from the UHRR comparison experiment. For each method and sample size setup (6 vs. 6, 12 vs. 12, 24 vs. 24, and 48 vs. 48) 20 simulations were performed with the following settings: normalization = ‘MR’, RNAseq = ‘bulk’, Protocol = ‘Read/UMI’, Distribution = ‘NB’, ngenes = 30000, nsims = 20, p.DE = 0.10. We verified with the data generated from the AML and NPC differentiation data that the gamma distribution (shape = 1, scale = 0.5) would be an appropriate log fold change distribution in this case (Figure S5B).

### Cell Preparation

Human embryonic kidney 293T (HEK293T) cells were cultured in DMEM media (TH.Geyer, L0102) supplemented with 10% FBS (Thermo Fisher, 10500-064) and 100 U/ml Penicillin and 100 μg/ml Streptomycin (Thermo Fisher). Cells were grown to 80% confluency, and harvested by trypsinization (Thermo Fisher, 25200072).

Peripheral blood mononuclear cells (PBMCs) were obtained from LGC Standards (PCS-800-011). Before use, the cells were thawed in a water bath at 37°C and washed twice with PBS (Sigma-Aldrich, D8537).

Prior to lysis, cells were stained with 1 ug/ml Trypan Blue (Thermo Fisher Scientific, 15-250-061) and counted using a Neubauer counting chamber. Then, the desired number of cells (1,000 or 10,000) was pelleted for 5 min at 200 rcf, resuspended in 50 µL of lysis buffer (RLT Plus (Qiagen, 1053393) and 1% ß-mercaptoethanol (Sigma-Aldrich,M3148) and transferred to a 96-well plate. Samples were then stored at -80 °C until needed.

### Tissue Preparation

Striatal tissue from C57BL/6 mice between the ages of 6 and 12 months was harvested by first placing the mouse in a container with Isoflurane (Abbot, TU 061220) until the mouse was visibly still and exhibited laboured breathing. The mice were then removed from the container, and a cervical dislocation was performed. The mice were briefly washed with 80% EtOH, the head decapitated, and the brain removed. The brain was transferred to a dish with ice-cold PBS and placed in a 1 mm slicing matrix.

Using steel blades (Wilkinson Sword, 19/03/2016DA), 5 coronal incisions were made. Biopsy punches (Kai Medical, BPP-20F) were then taken from the striatum and the tissue was transferred to a 1.5 mL tube with 50 µL of lysis buffer, RLT Plus and 1% ß-mercaptoethanol. The tubes were snap frozen and stored at -80 °C until needed.

### RNA Extraction Experiments

To determine differences due to RNA extraction we isolated RNA using columns from the Direct-zol RNA MicroPrep Kit (Zymo, R2062) (condition: “Column”) and magnetic beads from the prime-seq protocol (conditions: “No Incubation”, “Incubation”, and “Magnetic Beads”) (see above for details on prime-seq). For the “Column” condition, manufacturer instructions were followed and both the Proteinase K and DNase digestion steps were performed as outlined in the protocol. For the magnetic bead isolation, the prime-seq protocol was used as outlined in the “Magnetic Beads” condition. For “No Incubation” condition the Proteinase K digestion was skipped entirely. For the “Incubation” condition, the Proteinase K digestion was performed but with no enzyme; that is the heat cycling of 50°C for 15 minutes and 75°C for 10 minutes was carried out but no enzyme was added to the lysate.

### gDNA Priming Experiment

For a graphical overview of the gDNA Priming experiment, see Figure 2B. Frozen vials of mouse embryonic stem cells (mESC), that have been cultured as previously described (citation Bagnoli) (clone J1, frozen in Bambanker (NIPPON Genetics, BB01) on 04.2017), and HEK293T cells (frozen in Bambanker on 30.11.18, passage 25) were thawed. DNA was extracted from 1 million mESCs using DNeasy Blood & Tissue Kit (Qiagen, 69506) and RNA was extracted from 450,000 HEK293T cells using the Direct-zol RNA MicroPrep Kit (Zymo, R2062), according to manufacturer instructions in both cases. The optional DNase treatment step during the RNA extraction was performed in order to remove any residual DNA.

After isolating DNA and RNA, the two were mixed to obtain the following conditions: 10 ng RNA/ 7 ng DNA, 7.5 ng RNA/ 1.75 ng DNA, and 10 ng RNA/ 0 ng DNA. The 10 ng RNA/ 7 ng DNA condition, which represents the highest contamination of DNA, was performed twice, once without DNase treatment and once with DNase treatment. Libraries were prepared from three replicates for each condition using prime-seq and were then sequenced (see above for detailed information).

### MAQC-III Comparison Experiment

For a graphical overview of the experimental design see Figure S7A. As only Mix A from the original MAQC-III Study was compared, 122.2 µL of ddH2O, 2.8 µL of UHRR (100 ng/µL) (Thermo Fisher, QS0639), and 2.5 µL of ERCC Mix 1 (1:1000) (Thermo Fisher, 4456740) were combined to generate a 1:500 dilution of Mix A. Eight RNA-seq libraries were constructed using prime-seq (see above methods) with 5 µL of the 1:500 Mix A.

The samples were sequenced and the data processed and analyzed as outlined above. Of the comparison data from the original MAQC-III Study, Experiment SRX302130 to SRX302209 from Submission SRA090948 were used as this was the sequence data from one site (BGI) and was sequenced using an Illumina HiSeq 2000 [48]. The TruSeq data was first trimmed to be 50 bp long and then processed with zUMIs as outlined above, with the exception of using both cDNA reads and not providing UMIs as there were none. Paired-end data was used to not penalize TruSeq, as this is a feature of the method.

### NPC Differentiation Experiment

To differentiate hiPSCs to NPCs, cells were dissociated and 9×10^3^ cells were plated into each well of a low attachment U-bottom 96-well-plate in 8GMK medium consisting of GMEM (Thermo Fisher), 8% KSR (Thermo Fisher), 5.5 ml 100x NEAA (Thermo Fisher), 100mM Sodium Pyruvate (Thermo Fisher), 50mM 2-Mercaptoethanol (Thermo Fisher) supplemented with 500nM A-83-01 (Sigma Aldrich), 100nM LDN 193189 (Sigma Aldrich) and 30µM Y27632 (biozol). A half-medium change was performed on day 2 and 4. On day 6 Neurospheres from 3 columns were pooled, dissociated using Accumax (Sigma Aldrich) and seeded on Geltrex (Thermo Fisher) coated wells. After 2 days, cells were dissociated, counted and 2×10^4^ were lysed in 100 µL of lysis buffer (RLT Plus (Qiagen, 1053393) and 1% ß-mercaptoethanol (Sigma-Aldrich,M3148)

### AML-PDX Sample Collection

Acute myeloid leukemia (AML) cells were engrafted in NSG mice (The Jackson Laboratory, Bar Harbour, ME, USA) to establish patient derived xenograft (PDX) cells [52]. AML-PDX cells were cryopreserved as 10 Mio cells in 1mL of freezing medium (90% FBS, 10% DMSO) and stored at -80°C for biobanking purposes. To avoid thawing these samples and thus harming or even destroying them, the frozen cell stocks were first transferred to dry ice under a cell culture hood. Next a sterile 1 mm biopsy punch was used to punch the frozen cells in the vial and transfer the extracted cells to one well of a 96 well plate containing 100 µL RLTplus lysis buffer with 1% beta mercaptoethanol. To ensure complete lysis the lysate was mixed and snap frozen on dry ice. One biopsy punch is estimated to contain 10 µL of cryopreserved cells corresponding to roughly 1×10^5 cells given an even distribution of cells within the original vial. All 96 samples were collected in this manner, biopsy punches were washed using RNAse Away (Thermo Fisher Scientific) and 80 % Ethanol for reuse. These lysates were subjected to prime-seq, including RNA isolation using SPRI beads. In total, PDX samples from 11 different AML patients were analyzed in 6 to 16 biological replicates (engrafted mice) per sample.

### Cost Comparisons

Costs were determined by searching for general list prices from various vendors. When step by step protocols were available, each component was included in the cost calculation, such as for the SMARTer Stranded Total RNA Kit (Takara, 634862), SMART-Seq RNA Kit (v4) (Takara, 634891), TruSeq Library Prep (Illumina, RS-122-2001/2), TruSeq Stranded Library Prep (Illumina, 20020595), and Illumina Stranded mRNA Prep (Illumina, 20040534). In the case of BRB-seq no publicly available step-by-step protocol was found, so the methods section was used to calculate costs [22]. Decode-seq has a publicly available protocol, however, the level of detail was insufficient to calculate exact costs; therefore, when specific vendors were not listed, we used the most affordable option that we have previously validated. In all cases the prices included sales tax and were listed in euros and were therefore converted to USD using a conversion rate of 1.23 USD to EUR. The costs for all methods can be found in Table S4.

## Declarations

### Ethics approval and consent to participate

The human iPSC samples, which were differentiated into the NPCs, were ethically approved by the responsible committee on human experimentation (20-122, Ethikkommission LMU München) as previously published [57].

Bone marrow (BM) and peripheral blood (PB) samples from AML patients were obtained from the Department of Internal Medicine III, Ludwig-Maximilians-Universität, Munich, Germany. Specimens were collected for diagnostic purposes. Written informed consent was obtained from the patients. The study was performed in accordance with the ethical standards of the responsible committee on human experimentation (written approval by the Research Ethics Boards of the medical faculty of Ludwig-Maximilians-Universität, Munich, number 068-08 and 222-10) and with the Helsinki Declaration of 1975, as revised in 2013. All animal trials were performed in accordance with the current ethical standards of the official committee on animal experimentation (written approval by Regierung von Oberbayern; ROB-55.2Vet-2532.Vet_02-16-7 and ROB-55.2Vet-2532.Vet_03-16-56).

The mouse brain tissues were collected from mice that were bred and housed at the Biology Faculty Animal Facility at Ludwig Maximilian University in accordance with institutional ethical standards. The animal tissue was harvested according to the German Animal Welfare Act Paragraph 4 (organ removal for scientific reasons).

### Consent for publication

Not applicable

### Availability of data and materials

The datasets generated and/or analysed during the current study are available in the ArrayExpress repository under the following accession numbers E-MTAB-10133, 10138-10142, 10175. The code required to generate the figures can be found at https://github.com/Hellmann-Lab/prime-seq.

### Competing interests

The authors declare that they have no competing interests.

### Funding

This work was supported by the Deutsche Forschungsgemeinschaft (DFG) through the LMU Excellence Initiative, SFB1243 (Subproject A05/A14/A15), DFG EN 1093/2-1 (project number 406901759) and the Cyliax foundation.

### Authors’ contributions

AJ, LEW, CZ, and WE conceived the study. JG, AJ, and PN prepared iPSC, HEK293T, and tissue samples. JG performed differentiation experiments. BVick and IJ generated AML-PDX samples. DR and JWB designed the barcoded primers. AJ, LEW, JWB, and PN conducted the RNA-seq experiments. AJ and LEW performed sensitivity and gene expression analysis. LEW performed power analysis. BVieth and IH provided computational and statistical support. AJ, LEW, JWB, and WE wrote the manuscript. All authors read and approved the manuscript.

## Acknowledgements

We would like to thank Karin Bauer and Ming Zhao for lab support, Ines Bliesener and Maike Fritschle for animal work, Sabrina Schenk, Irena Stähler, and the staff at the LMU Biology Faculty Animal Facility for mouse colony maintenance, Dr. Stefan Krebs and the staff of LAFUGA for sequencing services, and Dr. Boyan Bonev and his lab for suggesting the Ultra II FS Kit as an alternative to tagmentation. Some illustrations in Figure 1, Figure 3 and Figure S2 were created with BioRender.com

## Supplementary Figures

**Figure S1.**
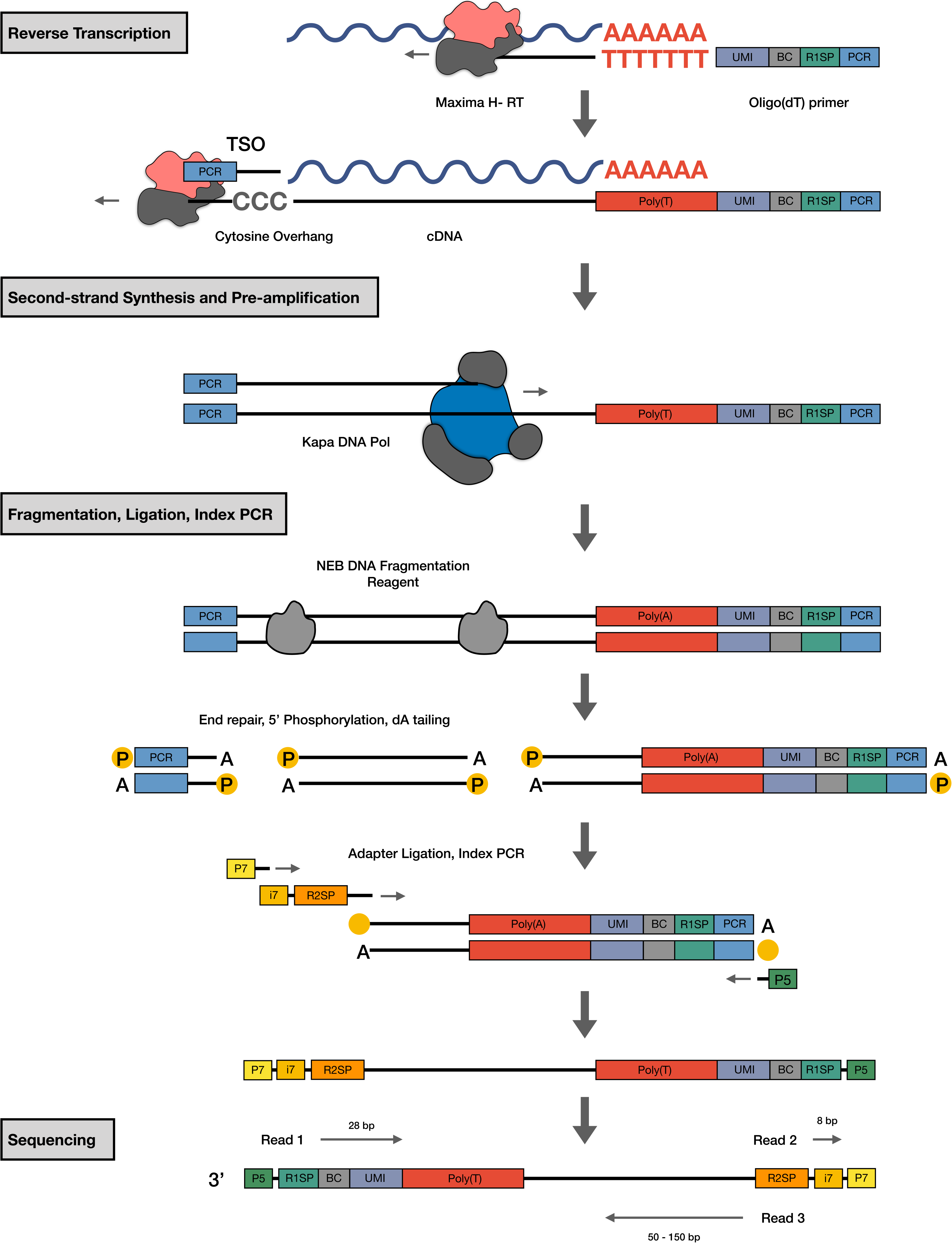
Molecular workflow of prime-seq. (Related to Figure 1) oligo(dT)-primers are used to enrich mRNA, which is then reverse transcribed using Maxima H-, a M-MLV reverse transcriptase. Full length first strand synthesis is performed using a template switching oligo. Second strand synthesis and cDNA pre-amplification is completed during the PCR using KAPA Hifi Polymerase, and this DNA is then used to generate libraries using the NEBNEXT Ultra II FS Kit. Finally the libraries are sequenced with the following setup: read 1: 28bp, read 2: 8bp, and read 3: 50-150bp.

**Figure S2.**
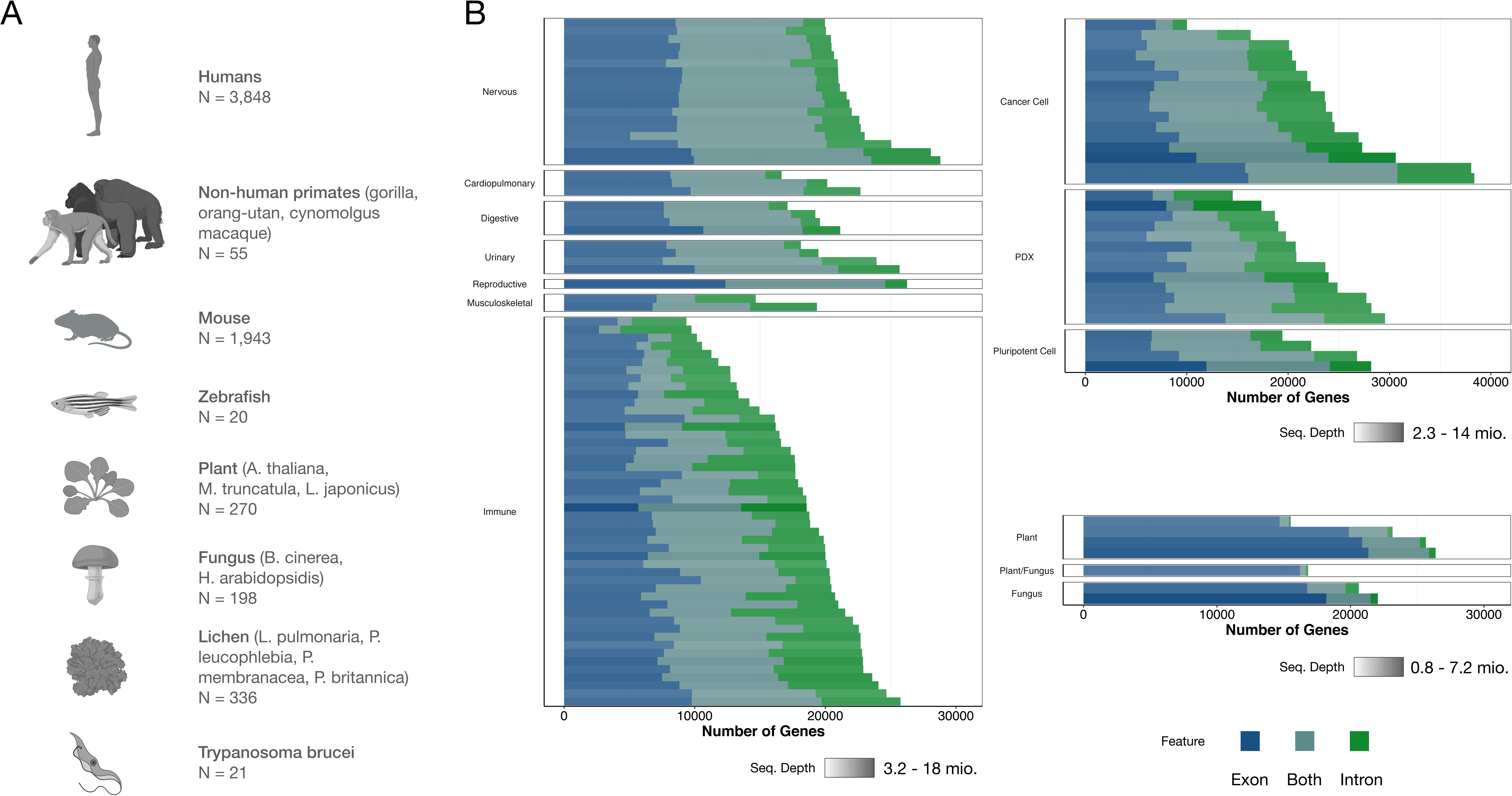
prime-seq is a robust protocol and has been validated with numerous organisms. (Related to Figure 2A) (A) To date, 132 experiments consisting of 6,691 samples from 17 different organisms, ranging from arabidopsis to zebrafish, have been processed with prime-seq. (B) Data from experiments with well-annotated genomes suggests a substantial number of detected genes come from intronic reads.

**Figure S3.**
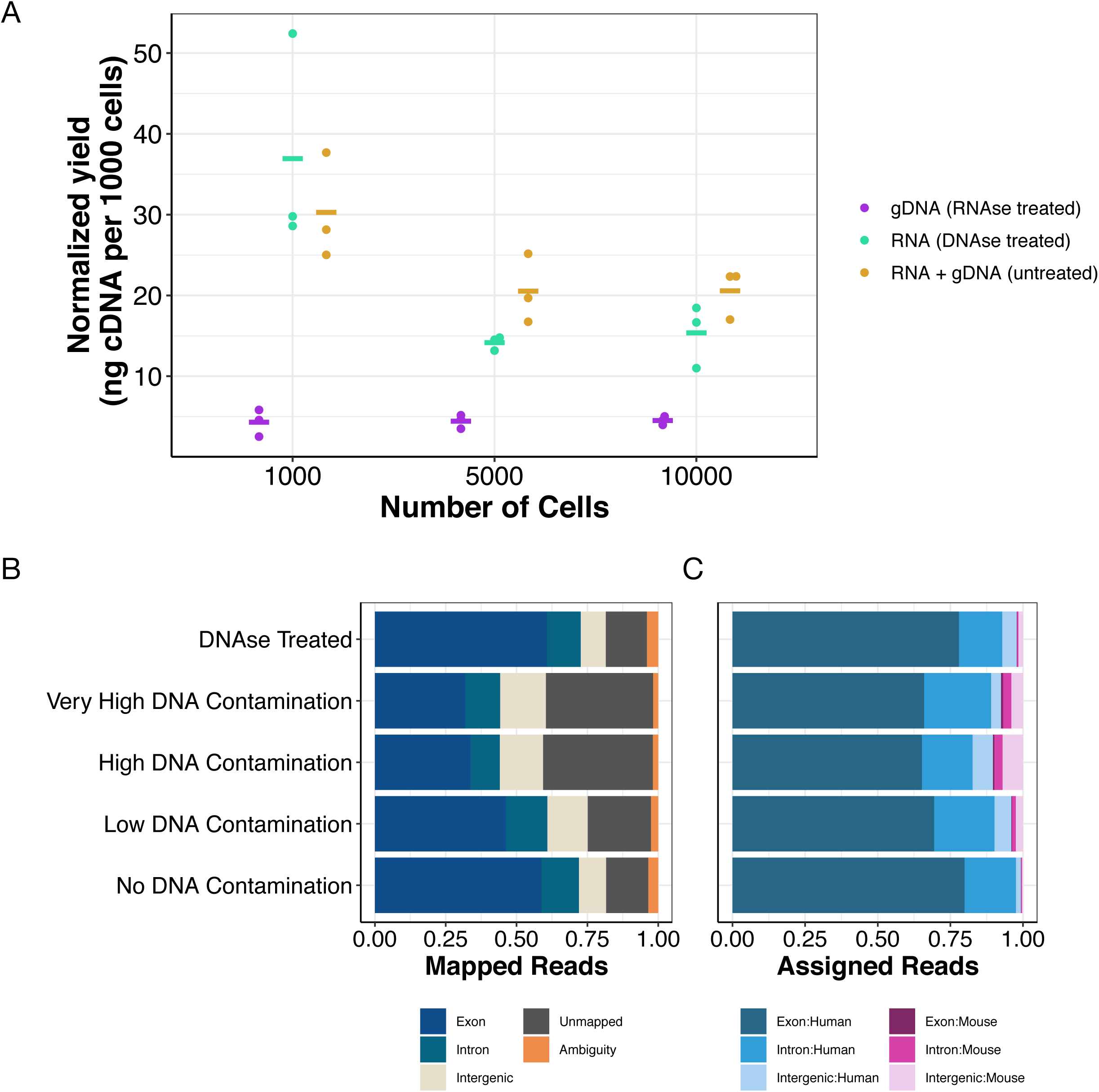
Intronic reads are not derived from contaminating gDNA. (A) Samples containing total nucleic acids were either treated with RNase A or DNase I, or remained untreated. Untreated samples had the highest concentration, showing that genomic DNA is also used as a template when not removed, albeit less efficiently than mRNA. cDNA yields were normalized to the number of input cells. (Related to Figure 2B) (B) Mapped reads from different gDNA/RNA mixed conditions, showing that the DNase treated condition and the no DNA contamination condition had the lowest fraction of intergenic and unmapped reads. (C) Fraction of assigned mapped reads per genomic feature (exon, intron, intergenic) and species, showing an increase in mouse reads with higher gDNA contamination.

**Figure S4.**
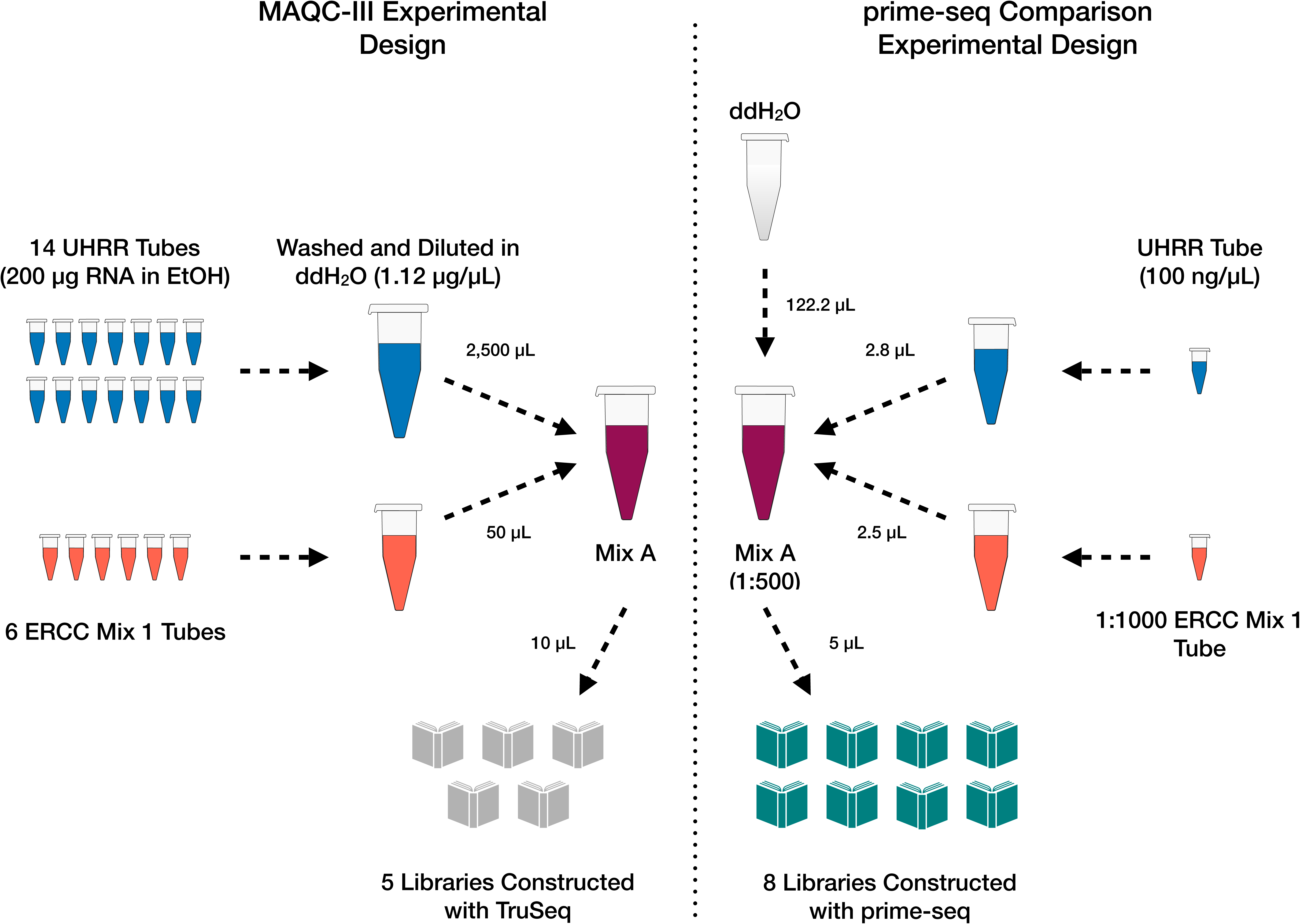
Experimental design comparing prime-seq to TruSeq data generated in the MAQC-III Study. (Related to Figure 3) A 1:1000 concentration of Mix A, from the MAQC-III Study, was generated by mixing UHRR and ERCC Mix 1. From this, eight libraries were generated using prime-seq and compared to five TruSeq generated libraries.

**Figure S5.**
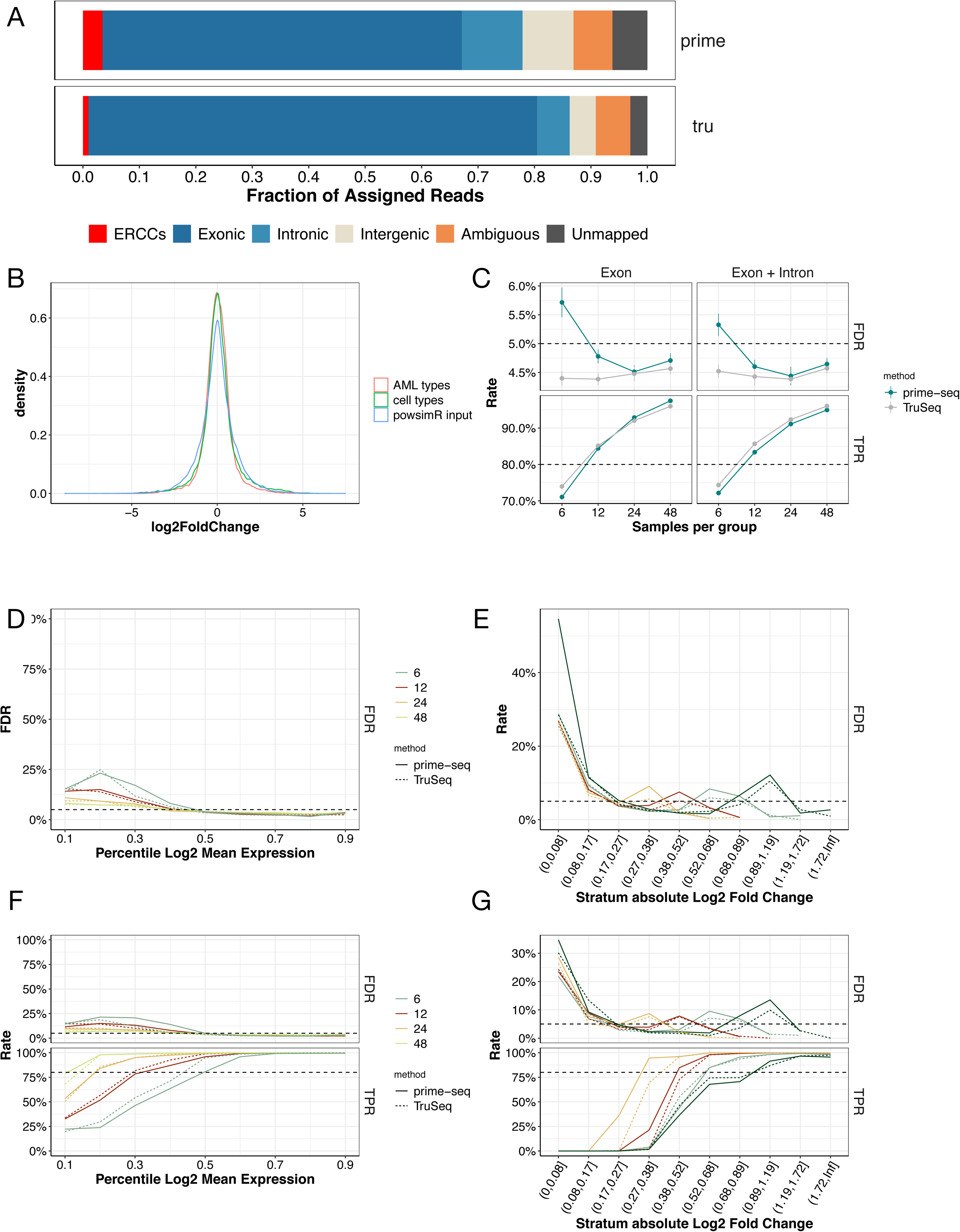
Power and FDR mostly depend on sample size and are similar between prime-seq and TruSeq. (Related to Figure 3) (A) Feature distribution from prime-seq and TruSeq shows 78% and 85% of reads are exonic, intronic, and ERCCs, respectively. (B) Log2 fold change distribution from the AML and NPC differentiation experiment (Figure 4) compared to the log2 fold change distribution used in powsimR for power analysis confirms that simulation settings match expected distributions. (C) Marginal power of prime-seq and TruSeq at differing samples per condition shows both methods perform similarly well, crossing the 80% threshold with roughly 12 samples both for exon plus intron and only exon counts. (D and E) FDR over different mean expression and log2 fold change strata (Related to 3G and 3H). (F and G) analogous to Figure 3G and 3H) but including only Exonic counts; prime-seq and TruSeq exhibit similar TPR and FDR over different mean expression and log2 fold change strata. Filtering parameters: detected UMI ≥ 1, detected gene present in at least 25%.

**Figure S6.**
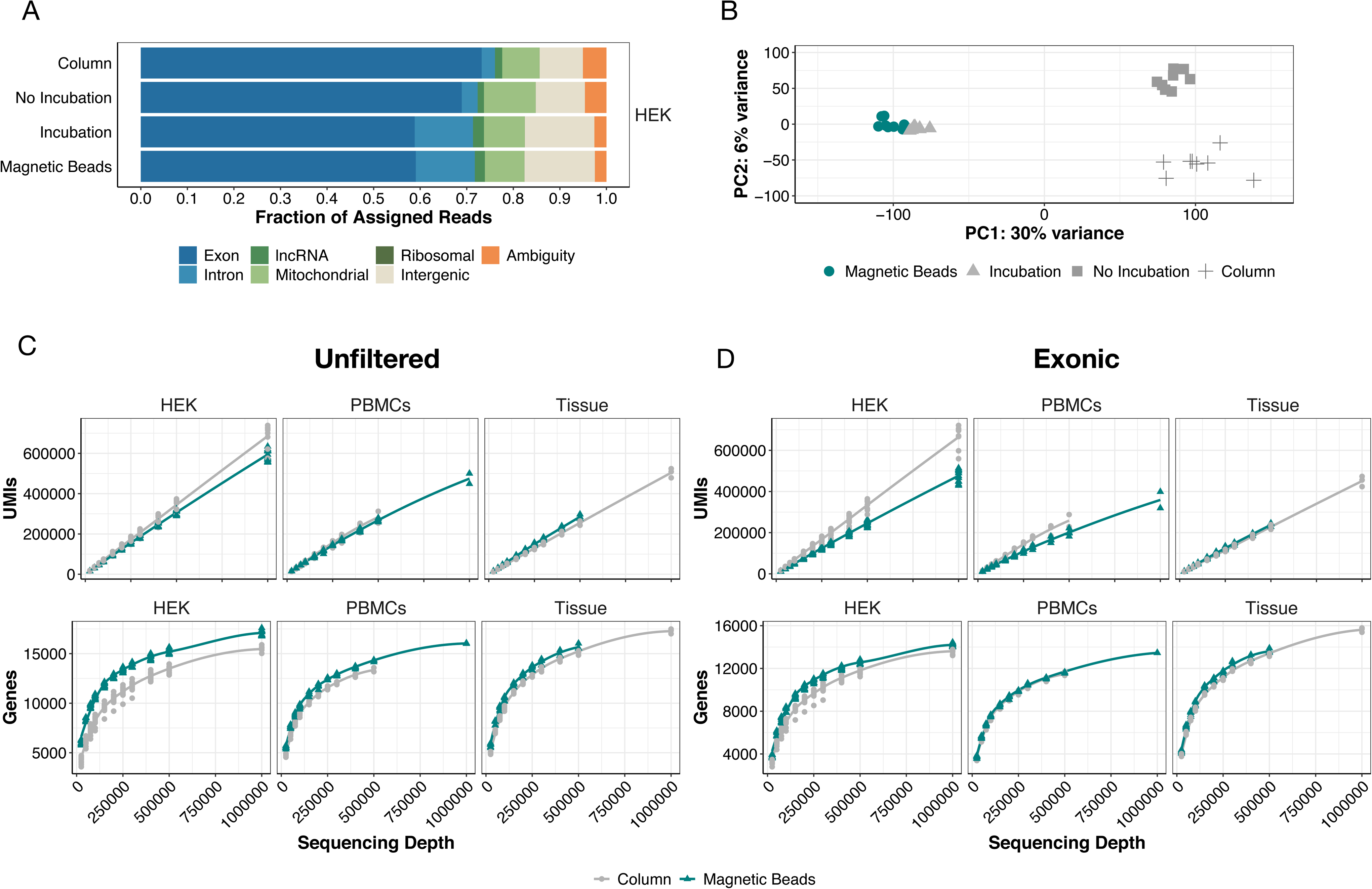
Performance of isolation methods is similar independent of prefiltering or usage of only Exon data. (Related to Figure 4) (A) HEK293T cell samples were extracted using columns and magnetic beads, employing the standard prime-seq protocol (“Magnetic Beads”), as well as variant protocols without proteinase K digestion (“No Incubation”) and a proteinase K digestion control without enzyme (“Incubation”). All conditions had similar fractions of usable reads (all but intergenic and ambiguity), with an increase in intronic reads in “Incubation” and “Magnetic Beads” suggesting this increase is due to heat incubation. (B) Principal component analysis (PCA) of the 500 most variable genes shows the largest variable is heat incubation. (C and D) Analysis of detected UMIs and detected genes for unfiltered data and exonic only data shows that prime-seq using magnetic bead isolation is more sensitive in HEK cells and similarly sensitive in PBMCs and tissue compared to prime-seq using column isolation. Filtering parameters: detected UMI ≥ 1, detected gene present in at least 25% of samples and is protein coding.

**Figure S7.**
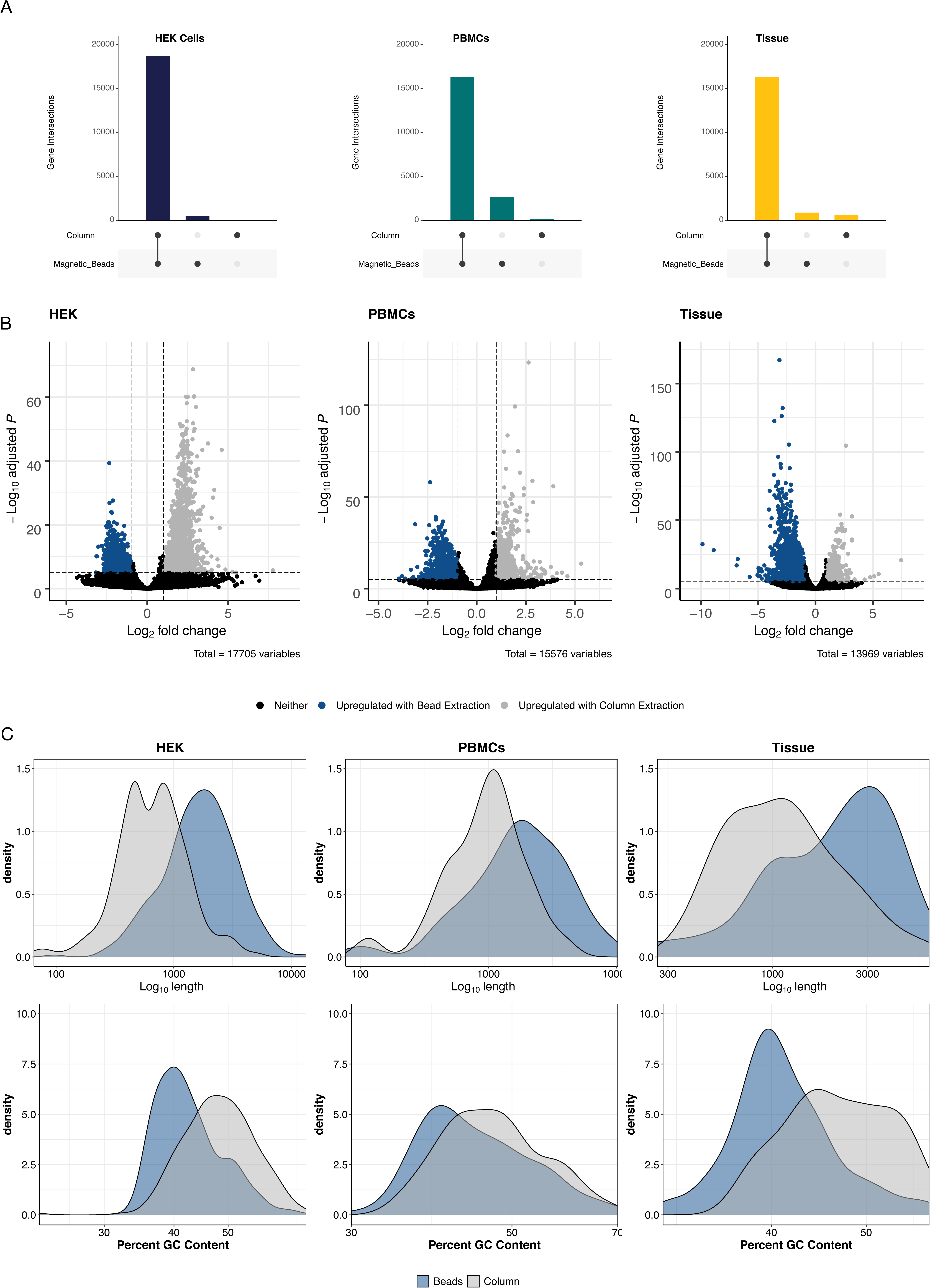
Most genes are detected independent of the extraction method used. (Related to Figure 4) (A) Upset plots showing a strong overlap of detected genes between columns and magnetic beads. (B) Up- and down-regulated genes between column and bead-based RNA extractions (p>0.05, log_2_ FC > 2). (C) Density plots of the differentially expressed genes relative to length and GC content. Genes upregulated in columns tend to be longer with lower GC content.

**Figure S8.**
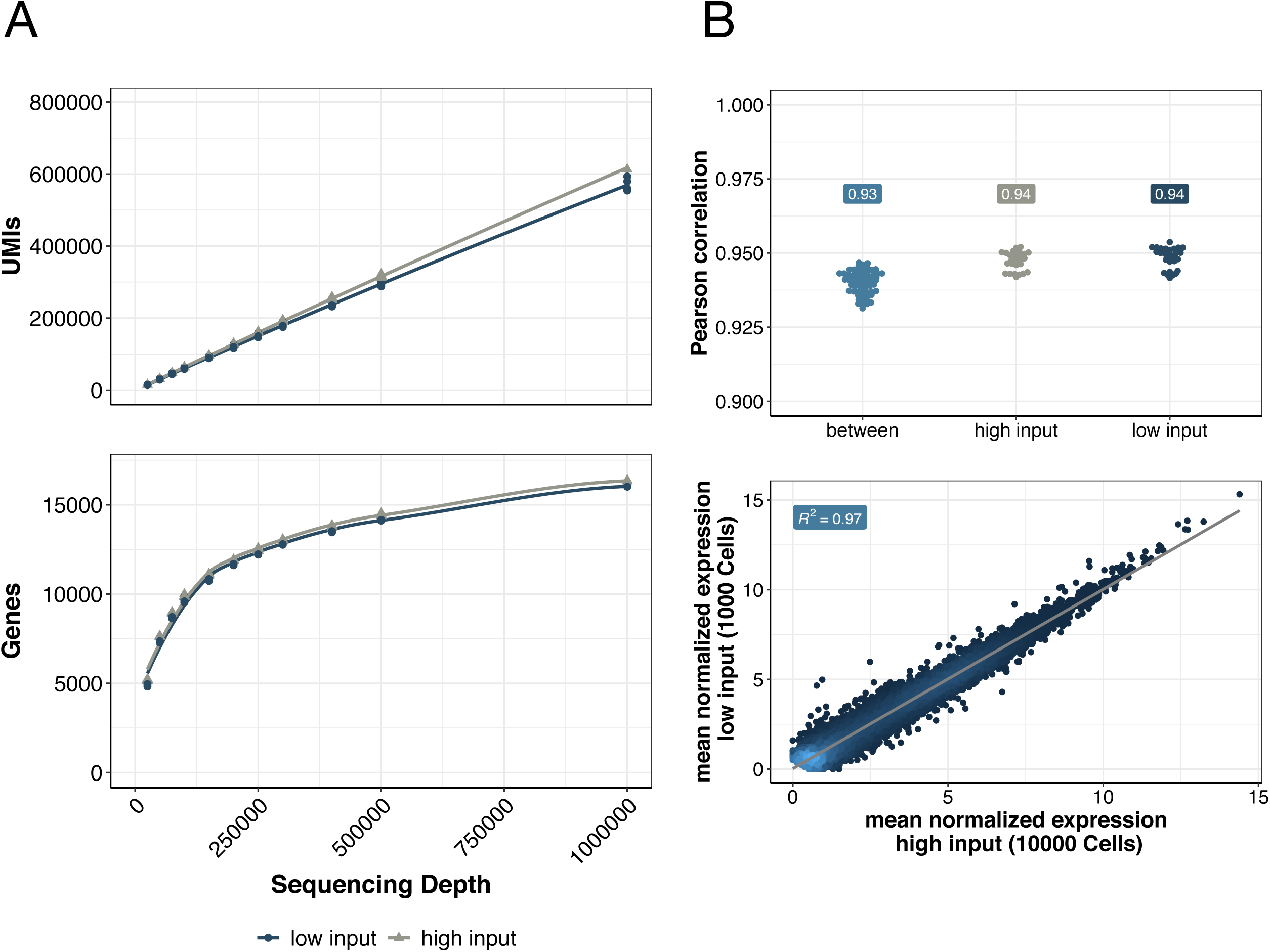
prime-seq performs equally well with high- and low-input samples. (Related to Figure 5) (A) Sensitivity, measured in detected UMIs and genes, is similar between high input (10,000 HEK293T cells) and low input (1,000 HEK293T cells) conditions at various sequencing depths (filtering parameters: detected UMI ≥ 1, detected gene present in at least 25% of samples and is protein coding). (B) Additionally, Pearson’s correlations between the high- and low-input conditions were high (pairwise comparison between: r = 0.93, pairwise comparison within: r = 0.94, and average normalized mean expression, R^2^ = 0.97).

**Figure S9.**
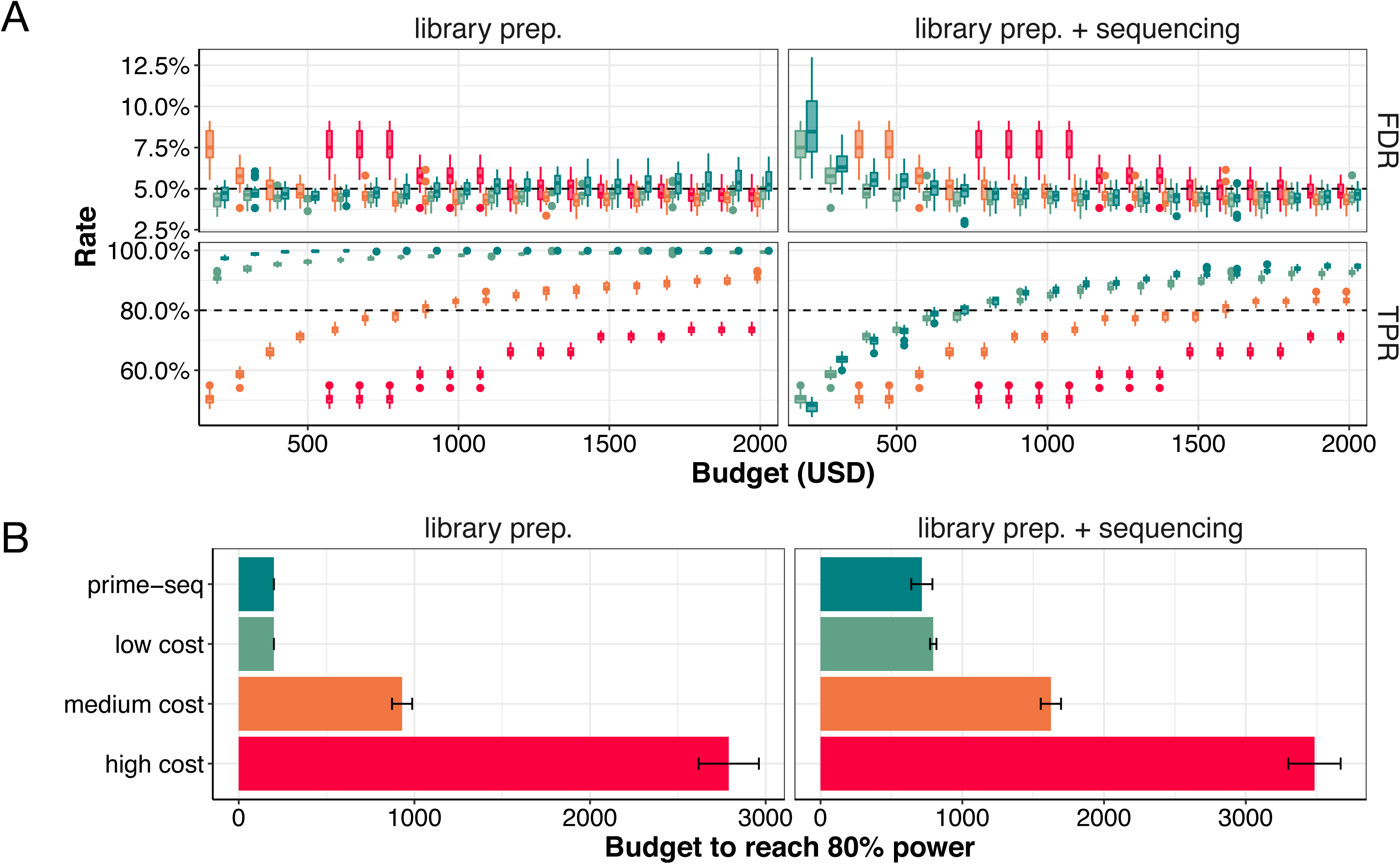
Power analysis shows prime-seq is able to reach 80% power earlier than less cost-efficient methods. (Related to Figure 6) (A) True positive rate (TPR) and false discovery rates (FDR) corresponding to Figure 6B, but with more incremental values. (B) prime-seq crosses an 80% power threshold with $715 when sequencing costs are included compared to $795, $1,625, and $3,485 for low, middle, and high cost methods respectively (10 million reads used for analysis at a cost of $3.40 per 1 mio. reads).

## Notes

### Competing Interest Statement

The authors have declared no competing interest.

## References

1. Stark R, Grzelak M, Hadfield J. RNA sequencing: the teenage years. Nat Rev Genet. 2019;20:631–56.

2. Ziegenhain C, Vieth B, Parekh S, Reinius B, Guillaumet-Adkins A, Smets M, et al. Comparative Analysis of Single-Cell RNA Sequencing Methods. Mol Cell. 2017;65:631–43.e4.

3. Vieth B, Parekh S, Ziegenhain C, Enard W, Hellmann I. A systematic evaluation of single cell RNA-seq analysis pipelines. Nat Commun. 2019;10:4667.

4. Svensson V, Vento-Tormo R, Teichmann SA. Exponential scaling of single-cell RNA-seq in the past decade. Nat Protoc. 2018;13:599–604.

5. Mereu E, Lafzi A, Moutinho C, Ziegenhain C, McCarthy DJ, Álvarez-Varela A, et al. Benchmarking single-cell RNA-sequencing protocols for cell atlas projects. Nat Biotechnol. 2020;38:747–55.

6. Ziegenhain C, Vieth B, Parekh S, Hellmann I, Enard W. Quantitative single-cell transcriptomics. Brief Funct Genomics. 2018;17:220–32.

7. Kivioja T, Vähärautio A, Karlsson K, Bonke M, Enge M, Linnarsson S, et al. Counting absolute numbers of molecules using unique molecular identifiers. Nat Methods. nature.com; 2011;9:72–4.

8. Hashimshony T, Wagner F, Sher N, Yanai I. CEL-Seq: single-cell RNA-Seq by multiplexed linear amplification. Cell Rep. 2012;2:666–73.

9. Parekh S, Ziegenhain C, Vieth B, Enard W, Hellmann I. The impact of amplification on differential expression analyses by RNA-seq. Sci Rep. 2016;6:25533.

10. Hagemann-Jensen M, Ziegenhain C, Chen P, Ramsköld D, Hendriks G-J, Larsson AJM, et al. Single-cell RNA counting at allele and isoform resolution using Smart-seq3. Nat Biotechnol. 2020;38:708–14.

11. Bagnoli JW, Ziegenhain C, Janjic A, Wange LE, Vieth B, Parekh S, et al. Sensitive and powerful single-cell RNA sequencing using mcSCRB-seq. Nat Commun. 2018;9:2937.

12. Zheng GXY, Terry JM, Belgrader P, Ryvkin P, Bent ZW, Wilson R, et al. Massively parallel digital transcriptional profiling of single cells. Nat Commun. 2017;8:14049.

13. Macosko EZ, Basu A, Satija R, Nemesh J, Shekhar K, Goldman M, et al. Highly Parallel Genome-wide Expression Profiling of Individual Cells Using Nanoliter Droplets. Cell. 2015;161:1202–14.

14. Klein AM, Mazutis L, Akartuna I, Tallapragada N, Veres A, Li V, et al. Droplet barcoding for single-cell transcriptomics applied to embryonic stem cells. Cell. 2015;161:1187–201.

15. Avila Cobos F, Alquicira-Hernandez J, Powell JE, Mestdagh P, De Preter K. Benchmarking of cell type deconvolution pipelines for transcriptomics data. Nat Commun. 2020;11:5650.

16. Li Y, Yang H, Zhang H, Liu Y, Shang H, Zhao H, et al. Decode-seq: a practical approach to improve differential gene expression analysis. Genome Biol. 2020;21:66.

17. Liu Y, Zhou J, White KP. RNA-seq differential expression studies: more sequence or more replication? Bioinformatics. 2014;30:301–4.

18. Lazic SE, Clarke-Williams CJ, Munafò MR. What exactly is “N” in cell culture and animal experiments? PLoS Biol. 2018;16:e2005282.

19. Subramanian A, Narayan R, Corsello SM, Peck DD, Natoli TE, Lu X, et al. A Next Generation Connectivity Map: L1000 Platform and the First 1,000,000 Profiles. Cell. 2017;171:1437–52.e17.

20. Uzbas F, Opperer F, Sönmezer C, Shaposhnikov D, Sass S, Krendl C, et al. BART-Seq: cost-effective massively parallelized targeted sequencing for genomics, transcriptomics, and single-cell analysis. Genome Biol. 2019;20:155.

21. Replogle JM, Norman TM, Xu A, Hussmann JA, Chen J, Zachery Cogan J, et al. Combinatorial single-cell CRISPR screens by direct guide RNA capture and targeted sequencing. Nat Biotechnol. Nature Publishing Group; 2020;38:954–61.

22. Alpern D, Gardeux V, Russeil J, Mangeat B, Meireles-Filho ACA, Breysse R, et al. BRB-seq: ultra-affordable high-throughput transcriptomics enabled by bulk RNA barcoding and sequencing. Genome Biol. 2019;20:71.

23. Ebinger S, Özdemir EZ, Ziegenhain C, Tiedt S, Castro Alves C, Grunert M, et al. Characterization of Rare, Dormant, and Therapy-Resistant Cells in Acute Lymphoblastic Leukemia. Cancer Cell. 2016;30:849–62.

24. Schreck C, Istvánffy R, Ziegenhain C, Sippenauer T, Ruf F, Henkel L, et al. Niche WNT5A regulates the actin cytoskeleton during regeneration of hematopoietic stem cells. J Exp Med. 2017;214:165–81.

25. Gegenfurtner FA, Zisis T, Al Danaf N, Schrimpf W, Kliesmete Z, Ziegenhain C, et al. Transcriptional effects of actin-binding compounds: the cytoplasm sets the tone. Cell Mol Life Sci. 2018;75:4539–55.

26. Gegenfurtner FA, Jahn B, Wagner H, Ziegenhain C, Enard W, Geistlinger L, et al. Micropatterning as a tool to identify regulatory triggers and kinetics of actin-mediated endothelial mechanosensing. J Cell Sci [Internet]. 2018;131. Available from: http://dx.doi.org/10.1242/jcs.212886

27. Mueller S, Engleitner T, Maresch R, Zukowska M, Lange S, Kaltenbacher T, et al. Evolutionary routes and KRAS dosage define pancreatic cancer phenotypes. Nature. 2018;554:62–8.

28. Wang S, Crevenna AH, Ugur I, Marion A, Antes I, Kazmaier U, et al. Actin stabilizing compounds show specific biological effects due to their binding mode. Sci Rep. 2019;9:9731.

29. Wang S, Gegenfurtner FA, Crevenna AH, Ziegenhain C, Kliesmete Z, Enard W, et al. Chivosazole A Modulates Protein-Protein Interactions of Actin. J Nat Prod. 2019;82:1961–70.

30. Ebinger S, Zeller C, Carlet M, Senft D, Bagnoli JW, Liu W-H, et al. Plasticity in growth behavior of patients’ acute myeloid leukemia stem cells growing in mice. Haematologica. 2020;105:2855–60.

31. Garz A-K, Wolf S, Grath S, Gaidzik V, Habringer S, Vick B, et al. Azacitidine combined with the selective FLT3 kinase inhibitor crenolanib disrupts stromal protection and inhibits expansion of residual leukemia-initiating cells in FLT3-ITD AML with concurrent epigenetic mutations. Oncotarget. 2017;8:108738–59.

32. Mulholland CB, Nishiyama A, Ryan J, Nakamura R, Yiğit M, Glück IM, et al. Recent evolution of a TET-controlled and DPPA3/STELLA-driven pathway of passive DNA demethylation in mammals. Nat Commun. 2020;11:5972.

33. Redondo Monte E, Wilding A, Leubolt G, Kerbs P, Bagnoli JW, Hartmann L, et al. ZBTB7A prevents RUNX1-RUNX1T1-dependent clonal expansion of human hematopoietic stem and progenitor cells. Oncogene. 2020;39:3195–205.

34. Shami A, Atzler D, Bosmans LA, Winkels H, Meiler S, Lacy M, et al. Glucocorticoid-induced tumour necrosis factor receptor family-related protein (GITR) drives atherosclerosis in mice and is associated with an unstable plaque phenotype and cerebrovascular events in humans. Eur Heart J. 2020;41:2938–48.

35. LaClair KD, Zhou Q, Michaelsen M, Wefers B, Brill MS, Janjic A, et al. Congenic expression of poly-GA but not poly-PR in mice triggers selective neuron loss and interferon responses found in C9orf72 ALS. Acta Neuropathol. 2020;140:121–42.

36. Geuder J, Ohnuki M, Wange LE, Janjic A, Bagnoli JW, Müller S, et al. A non-invasive method to generate induced pluripotent stem cells from primate urine [Internet]. Cold Spring Harbor Laboratory. 2020 [cited 2021 Jan 21]. p. 2020.08.12.247619. Available from: https://www.biorxiv.org/content/10.1101/2020.08.12.247619v1

37. Alterauge D, Bagnoli JW, Dahlström F, Bradford BM, Mabbott NA, Buch T, et al. Continued Bcl6 Expression Prevents the Transdifferentiation of Established Tfh Cells into Th1 Cells during Acute Viral Infection. Cell Rep. 2020;33:108232.

38. Kempf J, Knelles K, Hersbach BA, Petrik D, Riedemann T, Bednarova V, et al. Heterogeneity of neurons reprogrammed from spinal cord astrocytes by the proneural factors Ascl1 and Neurogenin2. Cell Rep. 2021;36:109409.

39. Porquier A, Tisserant C, Salinas F, Glassl C, Wange L, Enard W, et al. Retrotransposons as pathogenicity factors of the plant pathogenic fungus Botrytis cinerea. Genome Biol. BioMed Central; 2021;22:1–19.

40. Carlet M, Völse K, Vergalli J, Becker M, Herold T, Arner A, et al. In vivo inducible reverse genetics in patients’ tumors to identify individual therapeutic targets [Internet]. bioRxiv. 2020 [cited 2021 Sep 3]. p. 2020.05.02.073577. Available from: https://www.biorxiv.org/content/10.1101/2020.05.02.073577v1

41. Kempf JM, Weser S, Bartoschek MD, Metzeler KH, Vick B, Herold T, et al. Loss-of-function mutations in the histone methyltransferase EZH2 promote chemotherapy resistance in AML. Sci Rep. 2021;11:5838.

42. Pekayvaz K, Leunig A, Kaiser R, Brambs S, Joppich M, Janjic A, et al. Protective immune trajectories in early viral containment of non-pneumonic SARS-CoV-2 infection [Internet]. Cold Spring Harbor Laboratory. 2021 [cited 2021 Feb 19]. p. 2021.02.03.429351. Available from: https://www.biorxiv.org/content/10.1101/2021.02.03.429351v1

43. Kliesmete Z, Wange LE, Vieth B, Esgleas M, Radmer J, Hülsmann M, et al. TRNP1 sequence, function and regulation co-evolve with cortical folding in mammals [Internet]. Cold Spring Harbor Laboratory. 2021 [cited 2021 Feb 19]. p. 2021.02.05.429919. Available from: https://www.biorxiv.org/content/10.1101/2021.02.05.429919v2

44. Soumillon M, Cacchiarelli D, Semrau S, van Oudenaarden A, Mikkelsen TS. Characterization of directed differentiation by high-throughput single-cell RNA-Seq [Internet]. Cold Spring Harbor Laboratory. 2014 [cited 2021 Jan 21]. p. 003236. Available from: http://biorxiv.org/content/early/2014/03/05/003236.abstract

45. Parekh S, Ziegenhain C, Vieth B, Enard W, Hellmann I. zUMIs - A fast and flexible pipeline to process RNA sequencing data with UMIs. Gigascience [Internet]. 2018;7. Available from: http://dx.doi.org/10.1093/gigascience/giy059

46. Lee S, Zhang AY, Su S, Ng AP, Holik AZ, Asselin-Labat M-L, et al. Covering all your bases: incorporating intron signal from RNA-seq data. NAR Genom Bioinform [Internet]. Oxford Academic; 2020 [cited 2021 Jan 21];2. Available from: https://academic.oup.com/nargab/article-pdf/2/3/lqaa073/34054975/lqaa073.pdf

47. La Manno G, Soldatov R, Zeisel A, Braun E, Hochgerner H, Petukhov V, et al. RNA velocity of single cells. Nature. 2018;560:494–8.

48. Xu J, Su Z, Hong H, Thierry-Mieg J, Thierry-Mieg D, Kreil DP, et al. Cross-platform ultradeep transcriptomic profiling of human reference RNA samples by RNA-Seq. Sci Data. 2014;1:140020.

49. Vieth B, Ziegenhain C, Parekh S, Enard W, Hellmann I. powsimR: power analysis for bulk and single cell RNA-seq experiments. Bioinformatics. 2017;33:3486–8.

50. Oberacker P, Stepper P, Bond DM, Höhn S, Focken J, Meyer V, et al. Bio-On-Magnetic-Beads (BOMB): Open platform for high-throughput nucleic acid extraction and manipulation. PLoS Biol. 2019;17:e3000107.

51. Scholes AN, Lewis JA. Comparison of RNA isolation methods on RNA-Seq: implications for differential expression and meta-analyses. BMC Genomics. 2020;21:249.

52. Vick B, Rothenberg M, Sandhöfer N, Carlet M, Finkenzeller C, Krupka C, et al. An advanced preclinical mouse model for acute myeloid leukemia using patients’ cells of various genetic subgroups and in vivo bioluminescence imaging. PLoS One. 2015;10:e0120925.

53. Herold T, Jurinovic V, Batcha AMN, Bamopoulos SA, Rothenberg-Thurley M, Ksienzyk B, et al. A 29-gene and cytogenetic score for the prediction of resistance to induction treatment in acute myeloid leukemia. Haematologica. 2018;103:456–65.

54. Chambers SM, Fasano CA, Papapetrou EP, Tomishima M, Sadelain M, Studer L. Highly efficient neural conversion of human ES and iPS cells by dual inhibition of SMAD signaling. Nat Biotechnol. 2009;27:275–80.

55. Liu Y, Yu C, Daley TP, Wang F, Cao WS, Bhate S, et al. CRISPR Activation Screens Systematically Identify Factors that Drive Neuronal Fate and Reprogramming. Cell Stem Cell. 2018;23:758–71.e8.

56. Özdemir EZ, Ebinger S, Ziegenhain C, Enard W, Gires O, Schepers A, et al. Drug resistance and dormancy represent reversible characteristics in patients’ ALL cells growing in mice. Blood. American Society of Hematology; 2016;128:602–602.

57. Geuder J, Wange LE, Janjic A, Radmer J, Janssen P, Bagnoli JW, et al. A non-invasive method to generate induced pluripotent stem cells from primate urine. Sci Rep. 2021;11:3516.

58. Sholder G, Lanz TA, Moccia R, Quan J, Aparicio-Prat E, Stanton R, et al. 3’Pool-seq: an optimized cost-efficient and scalable method of whole-transcriptome gene expression profiling. BMC Genomics. 2020;21:64.

59. Ye C, Ho DJ, Neri M, Yang C, Kulkarni T, Randhawa R, et al. DRUG-seq for miniaturized high-throughput transcriptome profiling in drug discovery. Nat Commun. 2018;9:4307.

60. Pandey S, Takahama M, Gruenbaum A, Zewde M, Cheronis K, Chevrier N. A whole-tissue RNA-seq toolkit for organism-wide studies of gene expression with PME-seq. Nat Protoc. 2020;15:1459–83.

61. Kamitani M, Kashima M, Tezuka A, Nagano AJ. Lasy-Seq: a high-throughput library preparation method for RNA-Seq and its application in the analysis of plant responses to fluctuating temperatures. Sci Rep. 2019;9:7091.

62. Giraldez MD, Spengler RM, Etheridge A, Godoy PM, Barczak AJ, Srinivasan S, et al. Comprehensive multi-center assessment of small RNA-seq methods for quantitative miRNA profiling. Nat Biotechnol. 2018;36:746–57.

63. Xiong Y, Soumillon M, Wu J, Hansen J, Hu B, van Hasselt JGC, et al. A Comparison of mRNA Sequencing with Random Primed and 3’-Directed Libraries. Sci Rep. 2017;7:14626.

64. Picelli S, Björklund ÅK, Faridani OR, Sagasser S, Winberg G, Sandberg R. Smart-seq2 for sensitive full-length transcriptome profiling in single cells. Nat Methods. 2013;10:1096–8.

65. Westermann AJ, Vogel J. Cross-species RNA-seq for deciphering host-microbe interactions. Nat Rev Genet. 2021;22:361–78.

66. Dixit A. Correcting Chimeric Crosstalk in Single Cell RNA-seq Experiments [Internet]. bioRxiv. 2021 [cited 2021 Aug 26]. p. 093237. Available from: https://www.biorxiv.org/content/10.1101/093237v2

67. Gohl DM, Vangay P, Garbe J, MacLean A, Hauge A, Becker A, et al. Systematic improvement of amplicon marker gene methods for increased accuracy in microbiome studies. Nat Biotechnol. 2016;34:942–9.

68. Trück J, Eugster A, Barennes P, Tipton CM, Luning Prak ET, Bagnara D, et al. Biological controls for standardization and interpretation of adaptive immune receptor repertoire profiling. Elife [Internet]. 2021;10. Available from: http://dx.doi.org/10.7554/eLife.66274

69. Buschmann T, Bystrykh LV. Levenshtein error-correcting barcodes for multiplexed DNA sequencing. BMC Bioinformatics. 2013;14:272.

70. Somervuo P, Koskinen P, Mei P, Holm L, Auvinen P, Paulin L. BARCOSEL: a tool for selecting an optimal barcode set for high-throughput sequencing. BMC Bioinformatics. 2018;19:257.

71. Andrews S. FastQC: A quality control analysis tool for high throughput sequencing data [Internet]. Github; [cited 2021 Sep 14]. Available from: https://github.com/s-andrews/FastQC

72. Dobin A, Davis CA, Schlesinger F, Drenkow J, Zaleski C, Jha S, et al. STAR: ultrafast universal RNA-seq aligner. Bioinformatics. 2013;29:15–21.

73. Liao Y, Smyth GK, Shi W. The R package Rsubread is easier, faster, cheaper and better for alignment and quantification of RNA sequencing reads. Nucleic Acids Res. 2019;47:e47.

74. Team R. RStudio: Integrated Development for R. RStudio, PBC, Boston, MA, 2020. 2020.

75. R Core Team. R: A Language and Environment for Statistical Computing [Internet]. Vienna, Austria: R Foundation for Statistical Computing; 2016. Available from: https://www.r-project.org/

76. Steffen Durinck, Wolfgang Huber. biomaRt [Internet]. Bioconductor; 2017. Available from: https://bioconductor.org/packages/biomaRt

77. Wickham H, Francois R, Henry L, Müller K. dplyr: A grammar of data manipulation [Internet]. 2021. Available from: https://github.com/tidyverse/dplyr

78. Wickham H, Henry L. Tidyr: Tidy messy data. R package version. 2020;1:397.

79. Love MI, Huber W, Anders S. Moderated estimation of fold change and dispersion for RNA-seq data with DESeq2. Genome Biol. 2014;15:550.

80. Wickham H. ggplot2: Elegant Graphics for Data Analysis. Springer New York; 2010.

81. Wilke CO. cowplot: streamlined plot theme and plot annotations for “ggplot2.” 2019.

82. Clarke E, Sherrill-Mix S. ggbeeswarm: Categorical Scatter (Violin Point) Plots [Internet]. 2017. Available from: https://CRAN.R-project.org/package=ggbeeswarm

83. Constantin A-E, Patil I. ggsignif: R Package for Displaying Significance Brackets for “ggplot2” [Internet]. PsyArxiv. 2021. Available from: https://psyarxiv.com/7awm6

84. Xiao N. ggsci: Scientific Journal and Sci-Fi Themed Color Palettes for “ggplot2” [Internet]. 2018. Available from: https://CRAN.R-project.org/package=ggsci

85. Slowikowski K. ggrepel: Automatically position non-overlapping text labels with “ggplot2.” 2018.

86. Blighe K, Rana S, Lewis M. EnhancedVolcano: Publication-ready volcano plots with enhanced colouring and labeling. R package version. 2019;

87. Kremer LPM. ggpointdensity: A Cross Between a 2D Density Plot and a Scatter Plot [Internet]. 2019. Available from: https://CRAN.R-project.org/package=ggpointdensity

88. Kolde R. Pheatmap: pretty heatmaps [Internet]. 2012. Available from: https://cran.r-project.org/web/packages/pheatmap/index.html

